# Activation of Complement by Human Fibrin Clots: involvement of C1q and factor H

**DOI:** 10.1101/2023.10.19.563203

**Authors:** Yu-Hoi Kang, Praveen M. Varghese, Kirsten Pondman, Uday Kishore, Robert B. Sim

## Abstract

The classical pathway of the complement system is activated by the binding of C1q in the C1 complex to the target activator including immune complexes. Factor H is regarded as the key downregulatory protein of the alternative pathway of complement. However, C1q and factor H both bind to target surfaces via charge distribution patterns. For few targets, C1q and factor H compete for binding to common or overlapping sites. Factor H, therefore, can effectively regulate the classical pathway activation by such targets, in addition to its previously characterized role in the alternative pathway. Both C1q and factor H are reported to recognize “foreign” or altered-self materials. Clots, formed by the coagulation system, are an example of altered self. Factor H is present abundantly in platelets and is a well-known substrate for FXIIIa. Here, we investigated whether clots activate the classical pathway of complement and whether this is regulated by factor H. We show here that both C1q and factor H bind to fibrin formed in microtitre plates as well as fibrin clots formed under *in vitro* physiological conditions. Both C1q and factor H become covalently bound to fibrin clots and this is mediated via FXIIIa. We also show that fibrin clots activate the classical pathway of complement, as demonstrated by C4 consumption and membrane attack complex detection assays. Thus, factor H downregulates the classical pathway activation induced by fibrin clots. These results elucidate the intricate molecular mechanisms through which the complement and coagulation pathways intersect and have regulatory consequences.

## Introduction

The coagulation and the complement pathways are essential part of the innate immune system; they are known to be intertwined (1, 2) and thus influence each other’s initiation, effects and endpoints (3). The innate immune system is characterised by its ability to distinguish between “self”, “non-Self” and “altered self”. The complement system plays a crucial part in the innate immune surveillance. The complement system is activated via three pathways: Classical, Alternative, and Lectin (4).

C1q is the first recognition subcomponent in the complement classical complement (5). The binding of C1q to the target induces a conformational change in C1q which leads to activation of serine protease proenzyme C1r, which then activates proenzyme C1s, initiating the classical pathway. A variety of non-immunoglobulin activators have been shown to interact directly with C1q. These include nucleic acids and chromatin, mitochondrial membranes, possibly via cardiolipin or mitochondrial proteins, fibronectin, some viruses, Gram-positive bacteria via capsular polysaccharides, and Gram-negative bacteria via the lipid A component of the lipopolysaccharide (6). There is a considerable interest in the role of complement in clearing apoptotic cells, which has been suggested to occur by direct interaction of C1q with altered phospholipid distribution on these cells, including that of phosphatidylserine (7). Dysregulated activation of the complement system can cause permanent tissue or organ damage. Hence, the complement system is kept in check by various regulatory proteins to reduce undesired inflammatory responses and tissue damage. For example, factor H regulates the activation of the alternative pathway. Factor H binds soluble or membrane bound C3b and acts as a decay accelerating factor for C3bBb (8) or non-enzymatic cofactor for the cleavage of C3b to iC3b by factor I (9). C1q and factor H compete with each other for the same or overlapping binding sites, such as anionic phospholipids, lipid A and whole *E.coli* (10–13), suggesting that factor H is also able to act as a downregulator of the classical pathway (10–13).

The coagulation pathway is another cascade-driven homeostatic system (14). The coagulation system, composed of cells, proteins and processes that mediate blood clotting, is triggered by any damage to a blood vessel wall. This process is orchestrated through a series of proteolytic reactions (coagulation cascade), which is subdivided into the intrinsic, extrinsic, and common pathways (Graphic Abstract; Fig. 11). Thrombin, fibrinogen, and factor XIII (FXIII) play key roles in the last stages of the blood coagulation cascade. The central event in blood coagulation is to produce the coagulation enzyme, thrombin, which can convert soluble fibrinogen to insoluble fibrin (15, 16). The fibrinogen molecule is composed of two sets of three polypeptide chains termed α, β, γ, which are held together by disulfide bonds (17). This homodimer (αβγ)_2_ is an elongated 45-nm structure consisting of two outer D domains, each connected to a central E domain by a coiled-coil segment. E domain contains the fibrinopeptides A and B and the γ’ segment contains a thrombin and FXIII binding site (18). The serine protease thrombin cleaves the amino-terminal regions of the α and β chains of fibrinogen and releases 2 moles each of fibrinopeptide A (FPA) and fibrinopeptide B (FPB) for each mole of fibrin produced (19). Loss of FPA results in fibrin-I and additional loss of FPB yields fibrin-II. The fibrin II monomers polymerise through end-to-middle domain (D:E) association to form double-stranded fibrils, which then associate laterally to form fibrin fibres (20). These fibrin fibres form a network and the final fibrin solution is converted to a gel when at least 20 % of fibrinogen is converted to fibrin (18). Fibrin stabilization is accomplished by the action of factor XIIIa (plasma transglutaminase), formed by cleavage of soluble factor XIII by thrombin, which introduces numerous covalent crosslinks between these fibrin molecules. The resulting crosslinked fibrin web is able to capture platelets and red blood cells, effectively sealing the wound and stemming plasma loss. In addition to its primary role of providing scaffolding for the transvascular thrombus, fibrin participates in other biological functions involving unique binding sites (20). These include (i) suppression of plasma factor XIIIa-mediated crosslinking activity in blood by binding factor XIIIa; (ii) tissue type plasminogen activator (TPA) and plasminogen binding to fibrin which results in generation of plasmin, the major fibrinolytic protease (21); (iii) Leukocyte binding to fibrin(ogen) via integrin α_M_β_2_ (Mac-1) (22).

Plasma FXIII is a tetrameric molecule consisting of 2 A- and 2 B-polypeptides that are held together noncovalently (23). The A-subunit contains the enzyme active site and is synthesized by hepatocytes and monocytes. It contains an activation peptide of 37 amino acids that limits the access of the substrate to the active-site cysteine. The B-polypeptide serves as a carrier of the hydrophobic A-subunit in human plasma, is synthesized by the liver, and is secreted as a monomer that binds free A in plasma (24). Plasma FXIII is converted to the active FXIIIa in two steps. In the first step, thrombin cleaves an activation peptide from the A-subunit with formation of an inactive intermediate, FXIII’ (a’_2_b_2_) (25). A recent study showed that thrombin hydrolysis of the plasma FXIII activation peptides is accelerated in the presence of fibrin-I (26) through fibrin-I interaction with anion binding exosites-I (27) of FXIII. In the second step, calcium and fibrin induce the dissociation of the B-subunits from A to expose the active site thiol group. Fibrin polymers are an important cofactor to generate FXIIIa. The generation of FXIIIa in plasma can be triggered when approximately 1-2% of fibrinogen is converted to fibrin polymers (18). Activated FXIIIa first catalyses formation of γ-glutamyl-ε-lysyl bonds between fibrin γ-chains and then crosslinks the α-chains of fibrin monomers (28). In addition to being a critical component of the coagulation system, factor XIIIa also crosslinks fibronectin, vitronectin, collagen and lipoprotein in the extracellular matrix (29–32). Factor XIIIa also rapidly crosslinks α2-antiplasmin (an inhibitor of plasmin) to the α-chain of fibrin (33), which inhibits from the breakdown of fibrin by plasmin.

The complex interaction between the complement system and coagulation system is well-established (3). Briefly, anaphylatoxins C3a and C5a promote both inflammation and coagulation by activating platelets and inducing their aggregation. Activated platelets, which are fundamental constituents of the coagulation cascade, are implicated in the initiation of both the classical pathway and the alternative pathway of the complement system (34–36). Factor H has been localised to the α-granules of platelets and is released in response to thrombin stimulation or upon binding of C3b to the platelet surface (10, 37). Furthermore, factor H was found to copurify from platelets with thrombospondin-1 (38). Factor H is also a substrate for factor XIIIa (39–41). Thus, factor H present in platelets may participate in its interaction with the coagulation system as platelets may release factor H at the site of clotting, while FXIIIa may anchor factor H at the site. Factor H also inhibits FXI activation by thrombin or FXIIa (42). FXIa can cleave factor H reducing factor H binding to endothelial cells and thereby its activity in factor I mediated inactivation of C3b and C3b/Bb decay (43). Complement C1 inhibitor, a complement regulator, is another complement protein that can influence the coagulation process through inactivation of coagulation FXIIa (44). Likewise MASP-2, a critical enzyme in the complement lectin pathway, can cleave thrombin directly from prothrombin (45). The terminal complement component complex C5b–9 (membrane attack complex; MAC) also exhibits the ability to cleave prothrombin, even in the absence of Factor V (46). Cleaved complement component C5a, in particular, has been implicated in inducing procoagulant activity through a range of actions on endothelial cells and neutrophils, for example, by inducing an increase in the expression of Tissue Factor (TF) (47, 48) and instigating a switch in mast cell and basophil activities from a fibrin-dissolving (profibrinolytic) role to a clot-forming (prothrombotic) role via the upregulation of Plasminogen Activator Inhibitor-1 (PAI-1) (49). Similarly, thrombin promotes direct activation of C3 and C5, independent of conventional complement pathway activation (50). Thrombin and plasmin can activate complement during liver regeneration in the absence of C4 and alternative pathway activity (51). Several coagulation factors, specifically FIXa, FXa, FXIa, and plasmin, can directly cleave C3 and C5, resulting in the generation of potent anaphylatoxins C3a and C5a (52–54). FXIIa can also stimulate the classical pathway activation via the C1 complex (55, 56).

In this study, we examined the involvement of factor H with the coagulation system. We characterised factor H interaction with fibrin clots immobilised on microtiter wells and those formed more physiologically in human plasma. Activation of the classical pathway by fibrin clots, and the effects of factor H depletion on classical pathway activation were assessed.

## MATERIALS AND METHODS

### Fibrin clots

To prepare fibrinogen-coated wells, fibrinogen 5 μg/well (100 μl/well of 50 μg/ml) was incubated with 2 mM iodoacetamide and 1 mM Pefabloc-SC for 20 min at room temperature. The preincubated fibrinogen was then loaded onto microtiter plates (Maxisorp^TM^) and the plates were left for 1 h at 4°C. The plates were washed with PBS-0.5 mM EDTA, 0.1 % Tween 20 and blocked with the same buffer used in washing for 2 h at room temperature. To coat wells with fibrin, both fibrinogen (5 μg/well; 50 μl/well of 100 μg/ml) (without iodoacetamide and Pefabloc-SC) and thrombin (12.5 ng/well; 50 μl/well of 0.25 μg/ml) were diluted in 20 mM HEPES, 120 mM NaCl, 5 mM CaCl_2_, 0.05 mM DTT pH 7.4. Fibrinogen was dispensed onto microtitre wells and left for 10 min at room temperature. Then thrombin was added to the wells and left for 40 min at 37°C. Plates were then transferred to 4°C and left for 20 min. Plates were washed and blocked as for the fibrinogen-coated plates.

### Binding assays of C1q or factor H to fibrin- or fibrinogen-coated wells

C1q and factor H were purified from pooled human serum as described before (57, 58). C1q and factor H were radioiodinated as previously described (59, 60). ^125^I-C1q or ^125^I-factor H was serially diluted starting at 500,000 cpm/well (125 ng/well and 160 ng/well for C1q and factor H, respectively) in a volume of 100 μl of 25 mM HEPES-0.1% Tween 20 pH 7.4 or 100 μl of 25 mM HEPES–0.1 % Tween20, 5 mg/ml BSA pH 7.4. Dilutions were loaded onto fibrinogen- or fibrin- or BSA-coated wells and incubated for 30 min at 37°C. The wells were washed with HEPES buffer without BSA. To elute the bound ^125^I-C1q or ^125^I-factor H, 0.1 M NaOH was added to each well and allowed to incubate for 10 min. Supernatant was collected and counted in a Mini-Assay type 6-20 gamma counter (Mini-Instruments Ltd, Thermo Electronic Corporation, Reading, UK). This elution method was used in all following plate-binding assays.

To study saturation binding of ^125^I-factor H to fibrin, fibrin-coated wells were incubated with mixtures of a fixed amount of ^125^I-factor H (250,000 cpm, 80 ng/well) and various concentrations of unlabelled factor H (0-5120 ng/well) for 30 min at 37°C. Bound ^125^I-factor H was eluted and quantified as described above. The total amount of factor H bound was calculated from the radioactivity bound per well and the known total amounts of factor H supplied.

To determine the inhibitory effects of an excess of unlabelled factor H on binding of ^125^I-factor H to fibrin, ^125^I-factor H (2 μg/well) and different concentrations of unlabelled factor H (0-30 μg/well) or ovalbumin (0–9 μg/well) were premixed on ice, and then incubated with fibrin-coated wells for 30 min at 37°C. Bound ^125^I-factor H was eluted and quantified as described above.

To study the effect of ionic strength on the binding of ^125^I-C1q or ^125^I-factor H (100,000 cpm) to fibrin, fibrin-coated wells were incubated with ^125^I-factor H or ^125^I-C1q in each of four different veronal buffers (2.5 mM sodium barbital, 0.15 mM CaCl_2_, 0.5 mM MgCl_2_ pH 7.4) containing 0 mM, 75 mM, 150 mM or 500 mM NaCl for 30 min at 37°C. All washes were carried out at the same NaCl concentration that was used in the incubation. Bound ^125^I-factor H was eluted and quantified as described above.

### Binding assays of C1q or factor H to fibrin clots formed in human plasma

Binding of ^125^I-C1q and ^125^I-factor H to clots was examined at three different ionic strengths. Hence, plasma dialysed in HEPES-saline (90 %, final concentration) (0.5 ml) were diluted with (a) 0.5 ml water (to give final salt strength of about 70 mM); (b) 0.5 ml HEPES-saline (to give approximately 140 mM) or c) 1990 mM NaCl-10 mM HEPES-0.5 mM EDTA pH 7.4 (to give approximately 1 M). This gives a final concentration of 45 % plasma. A fixed amount of ^125^I-C1q or ^125^I-factor H (25,000 cpm) was premixed with each diluted plasma. CaCl_2_ (20 mM; final concentration) was then added and the mixtures were incubated for 40 min at 37°C. Clots were centrifuged (1,000 g, 5 min) at 4°C and supernatants were removed to measure radioactivity of unbound C1q or factor H. Clots were then washed three times with the appropriate reaction buffer and supernatants were removed. Radioactivity in the supernatants recovered from clotting reaction and washing was measured. ^125^I-C1q or ^125^I-factor H bound to fibrin clots was quantified by subtracting cpm remaining in the supernatants of washes from the initial cpm in each reaction. In order to examine if an increased clot size has an effect on C1q or factor H binding, experiments were modified by adding extra fibrinogen solution to a final concentration of 2 mg/ml in 100 μl of each reaction mixture. The clots formed in this way are called “enhanced clots”. The binding assays with additional fibrinogen added were carried out in the same manner as for non-enhanced clots. It was found that a high, easily measurable binding for both C1q and factor H was achieved at 70 mM salt strength with additional fibrinogen in clots so that enhanced clots formed at this salt strength was used in most experiments (unless indicated otherwise). Furthermore, in order to identify whether factor H and C1q bound directly to fibrin or to other proteins present in the plasma clots, it was necessary to examine the binding to fibrin clots which were made only from fibrinogen by treating with thrombin (denoted as fibrin-only-clots). Fibrin-only-clots were made from the same quantity of fibrinogen as present in the plasma for “enhanced clots”, in the presence of ^125^I-factor H or ^125^I-C1q.

To assess whether the binding of ^125^I-C1q or ^125^I-factor H to fibrin clots is covalent, fibrin clot-urea washing assay was performed. ^125^I-C1q or ^125^I-factor H (25,000 cpm) was incubated in clotting plasma for 8-time intervals in the range 0-960 min at 37°C. Clots were then washed 3 times vigorously with 500 μl of 10 mM HEPES, 70 mM NaCl, 0.5 mM EDTA, 5 M urea pH 7.4 and further unbound labelled protein was measured from the supernatants. ^125^I-C1q or ^125^I-factor H, which remained associated with clots was calculated as before. The remaining bound radioactive material is judged likely to be covalently bound. To investigate whether plasma proteins other than C1q and factor H interacted with fibrin clots, various radiolabelled plasma proteins were incubated in clotting plasma. The proteins used were ^125^I-C1q, ^125^I-factor H, ^125^I-human serum albumin (HSA), ^125^I-plasminogen, ^125^I-transferrin, ^125^I-IgG, ^125^I-α-2-Macroglobulin (25,000 cpm for each protein). HSA, plasminogen, IgG and α-2-Macroglobulin were purified at the MRC Immunochemistry Unit, Oxford and iodinated as described previously (59). Enhanced fibrin clots were formed in the presence of the plasma proteins. After 16 h incubation, fibrin clots were washed three times in 500 μl of 10 mM HEPES, 70 mM NaCl, 0.5 mM EDTA, 5 M urea pH 7.4. In order to show whether C1q or factor H was covalently linked to fibrin clots by the action of FXIIIa, ^125^I-C1q or ^125^I-factor H (25,000 cpm) was premixed with 10-fold diluted plasma (0.9%, final concentration in HEPES-½-saline) and 20 μg of fibrinogen in a total volume of 50 μl of HEPES-½-saline. CaCl_2_ was added to a final concentration of 20 mM and the mixture was incubated for 16 h at 37°C. As controls, either 20 mM ε-amino caproic acid (EACA) (final concentration) was added to mixtures before the addition of CaCl_2_ and preincubated for 5 min at room temperature, or 5 mM iodoacetamide (IAM) (final concentration) was added at the same stage for 10 min at room temperature. EACA and IAM are negative controls as they inhibit FXIIIa. Clots were washed three times with HEPES-½-saline or 10 mM HEPES, 70 mM NaCl, 0.5 mM EDTA, 5 M urea pH 7.4. Clots were then suspended in SDS-PAGE sample buffer containing 50 mM DTT and further analysed via SDS-PAGE (4-12% acrylamide) under reducing conditions.

### Haemolytic complement assay: Measurement of C4 consumption (via classical pathway)

Sensitized sheep erythrocytes (EA) for C4 consumption assay were prepared as described previously (8). Factor H-depleted plasma was prepared from normal human plasma as described earlier (61). In an ELISA, factor H-depleted plasma was then verified as being approximately 99.9% factor H deficient, compared to normal plasma (data not shown). C4 consumption by clot formation in normal human plasma and factor H-depleted plasma was determined by haemolytic assay. First, C4-deficient guinea pig serum (Sigma-Aldrich) was titrated on EA cells to select a suitable dilution, which did not cause complement-mediated lysis of the cells. Thus, C4-deficient guinea pig serum was serially diluted from 1/2 to 1/1024 in a volume of 100 μl with DGVB^2+^ in microtitre plates and then each dilution was incubated with 100 μl of EA cells in DGVB^2+^ (1 x 10^8^ cell/ml) for 1 h at 37°C. Cells were spun down at 1,000 g for 10 min and the optical density of the supernatants measured at 414nm. A positive control for 100% EA lysis was achieved through lysis of the erythrocytes with H_2_O. The negative control of DGVB^2+^ added to the EA cells gave the “0 % lysis” baseline. The background “0% lysis” was subtracted from each supernatant OD_414_ reading, and then the haemolytic activity was measured relative to 100% lysis positive control. As a result, 1/32 dilution of C4-deficient guinea pig serum was selected for use in assays to determine the changes in C4 levels (C4 consumption) by clots. Secondly, C4 consumption by fibrin clots was measured in normal human plasma (1/10 dilution in HEPES-½-saline) or factor H-depleted plasma (1/10 dilution in HEPES-½-saline) supplemented with fibrinogen (2 mg/ml) was allowed to clot in a total volume of 100 μl of HEPES-½-saline. Clotting was initiated by adding CaCl_2_ to 20 mM, and samples were incubated for 16 h at 37°C. Samples were centrifuged at 10,000g for 10 min at 4°C and supernatant was removed and further diluted 10-fold in DGVB^2+^.

This diluted supernatant (100 μl) was then loaded onto microtitre plates and serially diluted with DGVB^2+^ from 1/2 to 1/4096 (equivalent to final dilutions of the plasma of 1/200 to 1/409600). Each dilution was mixed with a fixed concentration of guinea pig serum (100 μl of 1/32 dilution in DGVB^2+^) on ice. EA cells (10^7^ cells/well in DGVB^2+^) were placed in microtitre plates and centrifuged for 10 min at 1,000g and the supernatants removed. The assay mixtures were then transferred to the wells containing EA cells and incubated for 1 h at 37°C. Cells were spun down at 1,000g for 10 min and OD_414_ of the supernatant was read. The percentage lysis was calculated as described above. C4 activity is expressed as the reciprocal dilution of serum/plasma required to give 50% lysis, it expressed as units of C4 activity per unit volume. The amounts of C4 consumption by clots were calculated by comparing with unclotted plasma. This assay measures the extent of C4 consumption (reduction in C4 level) occurring during clotting in the normal or factor H-depleted plasmas. Further control experiments were carried out, as above, with normal plasma and factor H-depleted plasma, but in the absence of clotting.

### Haemolytic complement assay: Measurement of MAC formation

MAC formation arising from complement activation by fibrin clots was determined by an ELISA system. MAC deposition was measured on fibrin-coated wells and MAC formation was also measured in the supernatants recovered from fibrin-coated wells incubated with serial dilutions of human serum. Ovalbumin antigen-antibody (OVA ag-ab) complexes and ovalbumin were used as a positive and a negative control, respectively. First, fibrin- or fibrinogen- (see above for preparation) or OVA ag-ab complexes- or ovalbumin-coated plates were prepared. OVA ag-ab complexes were prepared on plates as follows: hen ovalbumin (antigen) was dissolved in 0.1 M sodium carbonate pH 9.6. Ovalbumin (100 μl of 50 μg/ml) was added to each well and incubated for 1 h at room temperature. Plates were blocked with PBS-0.5 mM EDTA, 0.1 % Tween20 for 2 h at room temperature. Each well was then incubated with 200 μl of a 1:1 dilution of rabbit anti-ovalbumin antiserum (MRC Immunochemistry Unit, Oxford) in 1.5 M NaCl, 50 mM EDTA pH 7.4 for 1 h at room temperature. Plates were washed with 750 mM NaCl, 25 mM EDTA pH 7.4, and then washed again with PBS-0.5 mM EDTA, 0.1 % Tween 20. This gives OVA ag-ab complexes with some IgM but mostly IgG. The high NaCl concentration prevents binding of rabbit C1 or C1q. Secondly, fresh human serum was serially diluted (1/10 to 1/390625) in a total volume of 100 μl of DGVB^2+^ buffer on ice and immediately transferred to the fibrin-, fibrinogen-, ovalbumin-, OVA ag-ab complexes-coated wells and incubated for 1 h at 37°C. After incubation, plates were centrifuged at 1,000g for 10 min and supernatants were removed and kept at −20°C. The wells were washed with PBS-0.5 mM EDTA, 0.1 % Tween 20 and incubated with monoclonal anti-neo C9 antibody (3.75 μg/ml) for 1 h at room temperature. Mouse anti-neo C9 antibody (750 μg/ml in PBS-0.1% Azide) was a kind gift from Prof. Reinhard Wurzner (Innsbruck Medical University, Austria). It detects a neo epitope which is formed on C9 when incorporated into MAC. The wells were washed with PBS-0.5 mM EDTA, 0.1 % Tween20 and incubated with 100 μl of goat anti-mouse IgG alkaline phosphatase conjugate (Sigma) (1/5000 dilution in PBS-0.5 mM EDTA) for 1 h room temperature. The wells were washed again in PBS-0.5 mM EDTA, 0.1 % Tween 20 and visualised using the soluble alkaline phosphatase substrate p-nitrophenyl phosphate. Substrate solution (100 μl) was added to each well, incubated at room temperature in the dark and OD_405_ was read. Serial dilutions of fresh human serum were also incubated with blocked wells only to detect non-specific formation of MAC in the human serum and the values for these wells were subtracted from the values of each sample. Furthermore, MAC was also measured in the supernatants of serum incubated in wells using a capture ELISA system. In order to capture MAC in the supernatants, a capture antibody (rabbit anti-human C9 IgG) was purified from rabbit anti-human C9 antiserum using a HiTrap Protein G column (Pharmacia). Plates were coated with rabbit anti-human C9 IgG (5 μg/well in 0.1 M sodium carbonate pH 9.6) for 1 h at 20°C. After incubation, the wells were washed in PBS-0.5 mM EDTA, 0.1 % Tween 20 and blocked in the same buffer for 2 h at 20°C. The supernatants of serially diluted human serum removed from each fibrin-, fibrinogen, OVA ag-ab complexes-, ovalbumin-coated wells were then added to the wells and incubated for 1 h at room temperature. The wells were washed and MAC was detected using anti-neo C9 antibody, as described above. The supernatants of serially diluted fresh human serum recovered from blocked wells only were also assayed in the capture ELISA to detect background MAC formation in the human serum. Background values were subtracted from the values of each sample.

### Binding of ^125^I-C1q or ^125^I-factor H to “enhanced clots” formed in the presence or absence of plasma

Enhanced plasma clots were formed in the presence of ^125^I-C1q of ^125^I-factor H (25,000 cpm) as described above. Clots were washed with 500 μl of HEPES-1/2-saline buffer or 10mM HEPES, 70 mM NaCl, 0.5 mM EDTA, 5 M urea pH 7.4. The concentration of fibrinogen in the “enhanced clots” mixture is 3.5 mg/ml. Thus, to make clots in the absence of plasma, 3.5 mg/ml purified fibrinogen in HEPES-½-saline buffer was used. The fibrinogen solution (10 ml of 35 μg/ml) was premixed with ^125^I-C1q or ^125^I-factor H (25,000 cpm) in a total volume of 96.5 μl in 10 mM HEPES, 70 mM NaCl, 5 mM CaCl_2_ pH7.4. Clotting was initiated by adding 3.5 μl of thrombin (8.75 μg/ml, final concentration). This quantity of thrombin was selected as incubation of fibrinogen with thrombin at the same weight ratio resulted in complete cleavage of the α- and β-chains of fibrinogen at 40 min incubation at 37°C, as judged by our previous experiment (data not shown). As for the enhanced clots, the mixture was incubated for 40 min at 37°C and the resulting clots processed and washed as for the “enhanced clots”. ^125^I-C1q or ^125^I-factor H which remained associated with clots was measured as above.

### ELISA comparison of C1q or factor H in human plasmas and sera

C1q or factor H was detected from five different samples of citrated plasma, and the serum from the same plasmas. These plasmas represented a range of age in terms of storage and condition. They are Human Normal Plasma citrated (HPNC), Platelet-Poor Plasma (PPP Thawed), Pooled Citrated, Fresh plasma from a Healthy Volunteer (FHV) and O Rhesus Positive. HPNC, PPP Thawed and Pooled Citrated were all pooled outdated human plasma from HDS supplies. O Rhesus Positive was obtained from an eight-year old sample kept at 20°C. In order to detect C1q, OVA ag-ab complex-coated wells were used (see above for description). Serial two-fold dilutions of each plasma and serum pair from 1/10 to 1/5120 in PBS-0.1% Tween20-5 mM EDTA were incubated with the OVA ag-ab coated wells for 1 h at room temperature. The wells were washed with PBS-0.1% Tween 20 and then incubated with 100 μl of 100 μg/well of biotinylated rabbit anti-human C1q (Sigma) for 1 h at room temperature. The wells were washed and incubated with 100 μl/well of 1/40,000 extravidin alkaline phosphatase (Sigma) for 30 min at 20°C. The wells were washed and the plate was developed with p-nitrophenyl phosphate and OD was read at 405nm. Factor H was detected in the plasma and serum pairs as follows: monoclonal anti-human factor H MRCOX23 (MRC Immunochemistry Unit, 2.5 μg/well of purified IgG) coated-wells were used to capture factor H. Serial two-fold dilutions of each plasma and serum made from 1/20 to 1/10240 in PBS-0.1% Tween20 were incubated with the anti-factor H coated wells for 1 h at 20°C. The plates were washed with PBS-0.1% Tween20 and further incubated with rabbit anti-human factor H (MRC Immunochemistry Unit, Oxford) at a dilution of 1/32,000 in PBS-0.1% Tween20 for 1 h at 20°C. The wells were incubated with 100 μl of 1/10,000 dilution of goat anti-rabbit-IgG alkaline phosphatase conjugate (Sigma) for 1 h at 20°C. After incubation, the wells were washed and visualised using with alkaline phosphatase substrate, p-nitrophenyl phosphate (Sigma). The readings were taken at OD_405nm_.

## RESULTS

### Factor H and C1q interact with fibrin in the absence of BSA

Different concentrations of ^125^I-factor H or ^125^I-C1q were allowed to bind to fibrinogen-, fibrin-, or BSA-coated microtitre wells. Dose-dependent binding of ^125^I-factor H (Fig. 1A) or ^125^I-C1q (Fig. 1B) to fibrinogen or fibrin was observed. ^125^I-factor H and ^125^I-C1q both showed higher binding to fibrin than to fibrinogen although for C1q, the difference between fibrin and fibrinogen binding was small. When ^125^I-factor H (maximum quantity, 500,000, cpm/well) was added to fibrin-coated wells, approximately 1.7 % of factor H was bound. For ^125^I-C1q (500,000 cpm/well) approximately 2.4 % was bound to fibrin-coated wells. The binding of both ^125^I-factor H and ^125^I-C1q to fibrin was therefore low. ^125^I-C1q binding to fibrinogen- and fibrin-coated wells (only fibrin data are shown) was greatly decreased in the presence of fluid phase BSA, whereas binding of ^125^I-factor H was only slightly diminished by BSA (Fig. 1). ^125^I-factor H therefore, showed binding to fibrin which was “specific” in that it was not competed out by BSA. The amount of factor H bound was low but it was considered of sufficient interest to merit further investigation of the binding characteristics of ^125^I-factor H to fibrin. The binding of ^125^I-factor H and ^125^I-C1q to BSA-coated wells was very low. To determine the affinity of ligand binding to macromolecules (fibrin) and the number of receptors/fibrin-coated well, Scatchard plots were drawn where the X axis is specific binding (B) and the Y axis is specific binding divided by free radioligand concentration (B/F) as shown in Fig. 1 (top, left panel). The Scatchard plot for factor H binding was linear, suggesting that there was one class of receptors. The Kd (dissociation constant) indicates the strength of binding (affinity) between receptors and their ligands. If Kd is low, the affinity is high. For factor H Kd is 55.0 pM (slope equals −1/Kd where slope was −1.82 X10^−3^ fMol^−1^). This Kd value indicates exceptionally high affinity. The X-intercept was 20.5 fmol which is total ligand bound in fmol/well. Thus, each fibrin-coated well was able to bind to a maximum number of 1.234 × 10^10^ factor H molecules (maximum number of ligand molecule bound x Avogadro’s constant). That is, there are 1.234 x 10^10^ factor H binding sites/fibrin-coated well. Therefore, the affinity of factor H for fibrin is high, but number of ligand sites is low. The Scatchard plot representing ^125^I-C1q binding was non-linear. This is consistent with the multivalency of C1q.

**Figure 1:**
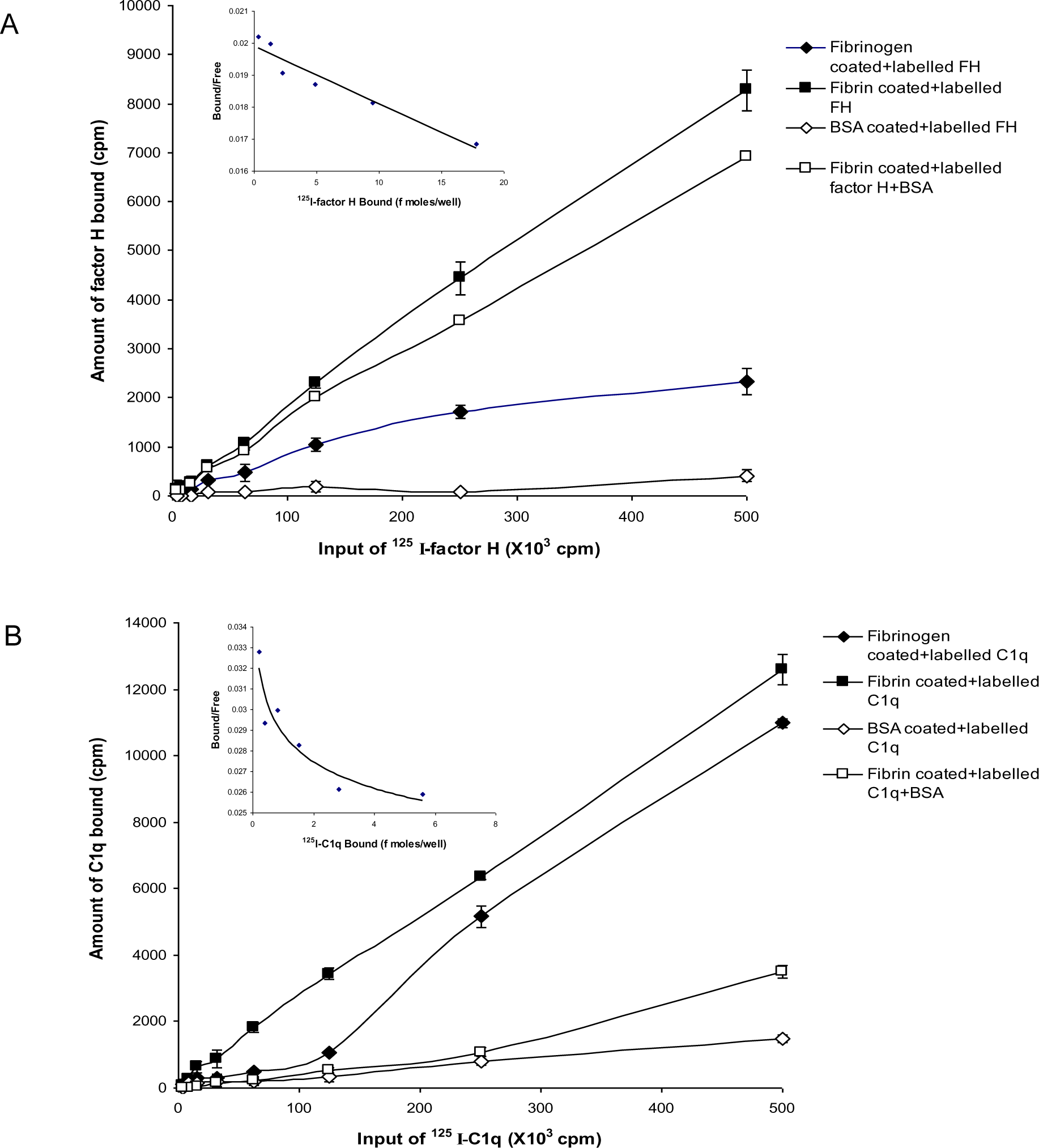
Binding of ^125^I-factor H, ^125^I-C1q to fibrinogen-, fibrin- and BSA-coated wells. A set of two-fold serial dilutions of ^125^I-factor H (A) or ^125^I-C1q (B) (starting at 500,000 cpm/well of ^125^I-factor H or ^125^I-C1q) were incubated with fibrinogen or fibrin for 30 min at 37°C in 20 mM HEPES-0.1% Tween20 pH 7.4. The wells were washed, and the amount of bound factor H and C1q were measured. BSA-coated wells were used as a negative control. Simultaneously, experiments were carried out with the sample dilutions of ^125^I-factor H or ^125^I-C1q in 20 mM HEPES-0.1% Tween 20 pH 7.4 containing 5 mg/ml BSA. All experiments were performed in duplicate and the average is shown. A Scatchard plot of Bound/Free versus Bound for each ^125^I-factor H or ^125^I-C1q binding to fibrin-coated wells are shown (*top, left panel*). The linear plot for factor H binding indicates that the fibrin clots have one receptor class (a single affinity receptor) with the slope of the line equal to −1/Kd. The intercept on the X-axis estimates the total ligand bound if all the receptor sites were occupied.

Binding of factor H to fibrin was dose-dependent and saturable. Saturation was achieved at an input of approximately 3.5 μg of factor H (Fig. 2A). Approximately 3.5 ng of factor H was bound to fibrin-coated wells with the input of 3.5 μg, thus 0.1% of binding was achieved. To determine the inhibitory effects of excess of unlabelled factor H on binding of ^125^I-factor H to fibrin, both labelled factor H (2 μg) and unlabelled factor H (0-30 μg/well) were premixed on ice then incubated with fibrin-coated wells. The results showed that there was a decrease in ^125^I-factor H binding to fibrin-coated wells as the concentration of unlabelled factor H increases (Fig. 2B). At an equal molar ratio of unlabelled to labelled ligand, unlabelled factor H inhibited binding of ^125^I-factor H to fibrin-coated wells by 48 %. Therefore, unlabelled factor H predictably inhibits labelled factor H binding. Ovalbumin did not interfere with ^125^I-factor H binding to fibrin (Fig. 2B).

**Figure 2.**
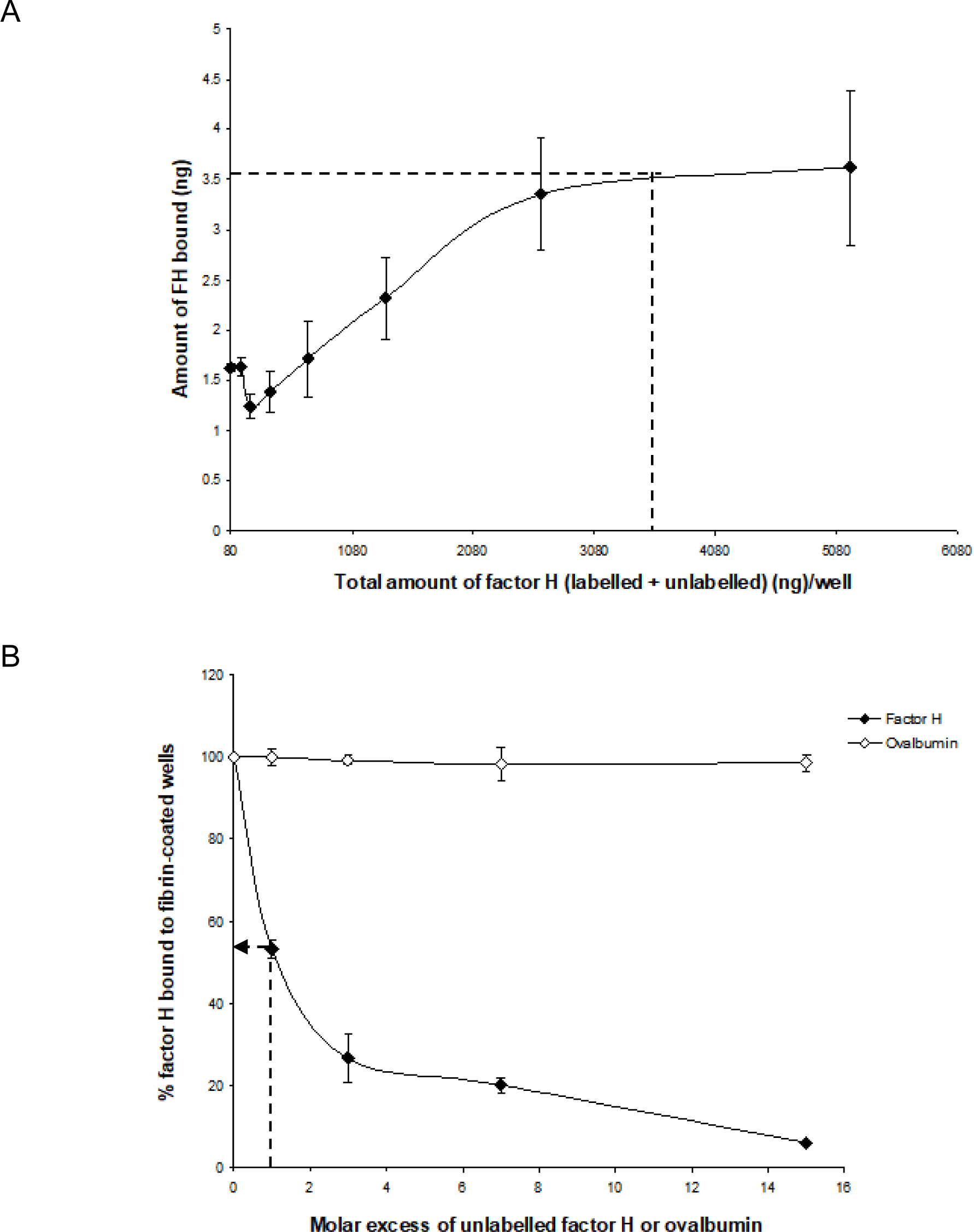
Characterisation of binding of ^125^I-factor H to fibrin-coated wells. Saturation of binding of ^125^I-factor H to fibrin-coated wells (A). Various amounts of unlabelled factor H (0–5120 ng/well) was mixed with a fixed amount of ^125^I-factor H (250,000 cpm/well equivalent to 80 ng/well) on ice and loaded on to fibrin-coated wells. Saturation was observed at an input of approximately 3.5 μg of factor H per well (denoted by a dotted mark). The means of three experiments are presented with standard deviations. Inhibition of ^125^I-factor H binding to fibrin by excess factor H (B). Labelled factor H (2 μg/well) was incubated with increasing amounts of unlabelled factor H (0–30 μg/well) in fibrin-coated wells for 30 min at 37°C. Wells were washed and the amount of ^125^I-factor H bound was measured as cpm bound per well. Ovalbumin (0-9 mg/well) was used as a control. The dotted arrow represents the percentage of factor H bound to fibrin when a 1:1 molar ratio of factor H to C1q was used. The means of three experiments are presented with standard deviations.

To determine whether the interaction of ^125^I-factor H or ^125^I-C1q with fibrin-coated wells is affected by ionic strength, ^125^I-factor H or ^125^I-C1q in veronal buffer with four different ionic strengths (20 mM NaCl, 75 mM NaCl, 150 mM NaCl, and 500 mM NaCl) were incubated in fibrin-coated wells. Very high binding of ^125^I-factor H or ^125^I-C1q was observed at 20 mM NaCl (Fig. 3). Binding at 75 mM NaCl concentration was less than 50 % of that seen at 20 mM NaCl. The binding of ^125^I-factor H and ^125^I-C1q was greatly decreased at 150 mM NaCl. At 500 mM NaCl, binding was not detectable for either ^125^I-factor H or ^125^I-C1q. This shows that binding of ^125^I-factor H or ^125^I-C1q is ionic in nature. Furthermore, both factor H and C1q binding was independent of divalent metal ions as a similar binding was observed with HEPES-saline-EDTA (5 mM EDTA was added to HEPES-saline) and HEPES-½-saline (data not shown).

**Figure 3.**
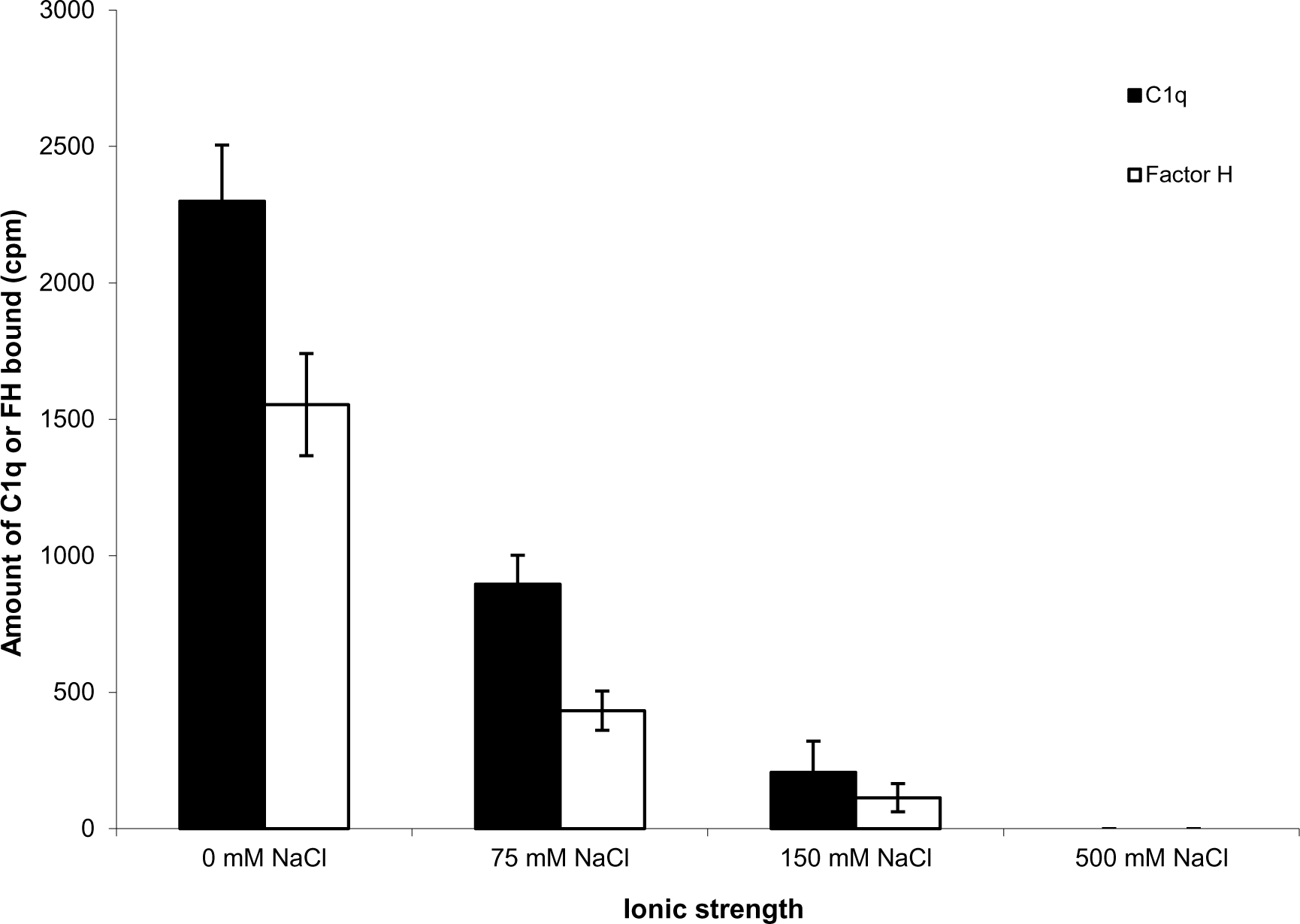
Effect of ionic strength on binding of ^125^I-factor H or ^125^I-C1q to fibrin. ^125^I-factor H (32 ng; 100,000 cpm) or ^125^I-C1q (20 ng; 100,000 cpm) at four different ionic strengths (20 mM, 75 mM NaCl, 150 mM NaCl and 500 mM NaCl) in VB^2+^ buffer was incubated with fibrin-coated wells. The means of three experiments are presented with standard deviations.

### ^125^I-factor H and ^125^I-C1q interact with clots formed in plasma

Factor H and probably C1q seem to bind to fibrin-coated wells as determined by plate assays (Fig. 1-3). It was therefore of interest to examine whether an interaction of ^125^I-factor H or ^125^I-C1q with fibrin clots can be measured in more physiological conditions. The coating of plates with fibrin/fibrinogen used in this work is quite artificial as the plate-fixed fibrin cannot move to form a fibrin polymer as occurs in clots. To get closer to “real” conditions, fibrin clots were formed in plasma in the presence of ^125^I-factor H or ^125^I-C1q. Before studying the binding of factor H or C1q to fibrin clots, the quantity of ^125^I-fibrinogen incorporated into crosslinked fibrin clots was determined (data not shown). Also, the efficiency of washing of fibrin clots was examined. Approximately 89 % of ^125^I-fibrinogen was incorporated into clots. Unbound ^125^I-fibrinogen or fibrin (approximately 8.5 %) was removed mostly in the first supernatant of the clotting reaction and only a small amount of ^125^I-fibrinogen or fibrin (< 1%) was detected in all three washes. Unbound ^125^I-fibrinogen may occur because not all of the fibrinogen is activated by thrombin, or not all of the fibrin is crosslinked by FXIIIa.

Binding of ^125^I-factor H and ^125^I-C1q to clots was examined at three different ionic strengths. Clots were formed in human plasma in the presence of ^125^I-factor H or ^125^I-C1q at three different salt concentrations and the percentage of ^125^I-factor H or ^125^I-C1q associated with the clot was determined by measuring depletion of radioactivity from the supernatant after clot formation. The binding of ^125^I-factor H and ^125^I-C1q were 21.5% and 2.5%, respectively at 70 mM salt concentration (Fig. 4). There was also some detectable binding at 1 M NaCl. The percentage binding of both factor H and C1q to these fibrin clots was much greater than that observed in plate assays. In order to optimise clot formation and potential binding to clots by ^125^I-factor H or ^125^I-C1q, fibrinogen (2 mg/ml, final concentration) was added to the reaction mixture prior to clot formation. This produces “enhanced” fibrin clots. In comparison to the binding shown in Fig. 1, there was “enhanced” binding of ^125^I-factor H and ^125^I-C1q observed at several different salt concentrations when fibrinogen was added in order to increase the clot size (Fig. 4B). The percentage of bound ^125^I-factor H was increased 6-fold at physiological salt concentration (140 mM NaCl) but no significant increase in binding was observed at 70 mM salt concentration. ^125^I-C1q binding was enhanced compared to normal clots at all salt concentrations. Therefore, the binding of ^125^I-factor H and ^125^I-C1q to fibrin was greater when clot volume/surface area was increased by adding more fibrinogen into the clotting mixture. This will increase the concentration of “epitopes” for factor H or C1q binding. Although increasing salt strength may have altered the kinetics of clot formation, clot formation was observed by eye at all the strengths tested.

**Figure 4.**
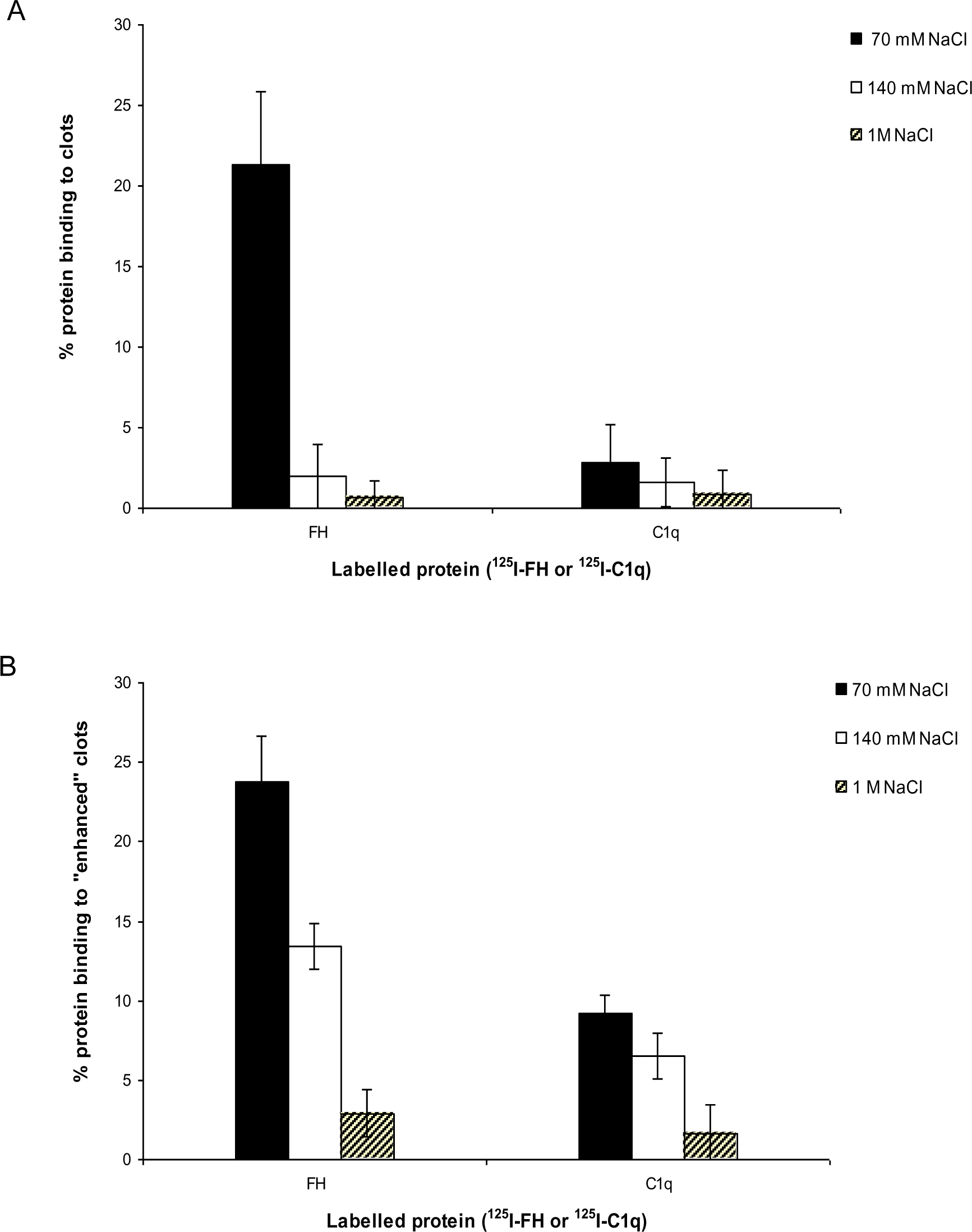
Interaction of ^125^I-factor H or ^125^I-C1q with plasma clots formed in more physiological condition. (A) Clots were formed in plasma in the presence of ^125^I-factor H (5 ng, 25,000 cpm/reaction) or ^125^I-C1q (8 ng, 25,000 cpm/reaction) at three salt strengths (70 mM, 140 mM and 1 M NaCl). (B) “Enhanced” clots were also formed in the presence of ^125^I-factor H or ^125^I-C1q (25,000 cpm/reaction) at different ionic concentrations (70 mM, 140 mM, 1 M NaCl). The means of three experiments are presented with standard deviations.

### Evidence for covalent binding of ^125^I-factor H and ^125^I-C1q to enhanced fibrin clots

Further experiments were carried out with enhanced fibrin clots. To see if the binding of ^125^I-factor H and ^125^I-C1q to fibrin clots is covalent i.e. involving crosslinking by FXIIIa, a kinetic experiment of ^125^I-factor H or ^125^I-C1q binding to fibrin clots was performed. After incubation for various times, fibrin clots with bound ^125^I-factor H or ^125^I-C1q were washed in 10 mM HEPES, 70 mM NaCl, 0.5 mM EDTA, 5 M urea pH 7.4 in order to denature non-covalently bound proteins. The proportions of ^125^I-factor H and ^125^I-C1q binding to clots increased as incubation time increased (Fig. 5). The binding of both ^125^I-factor H and ^125^I-C1q reached its maximum level after 100-180 min incubation as assessed after washing with HEPES-½-saline.

**Figure 5.**
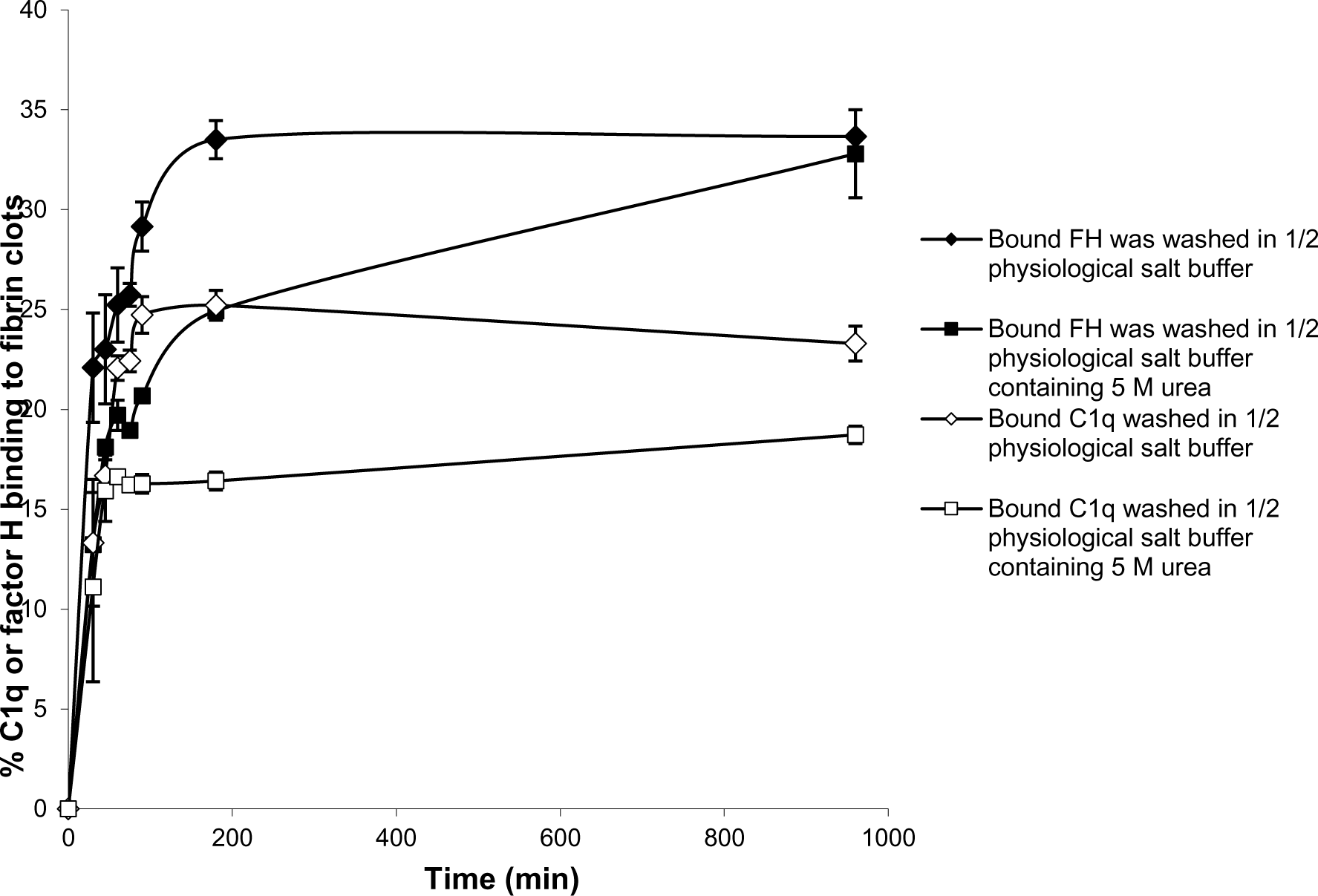
Time course of ^125^I-factor H and ^125^I-C1q binding to fibrin clots. Enhanced fibrin clots were formed in the presence of ^125^I-factor H or ^125^I-C1q, but the clotting mixtures were incubated for 8 time intervals in the range 0-960 min at 37°C. The enhanced clots were washed in HEPES-½-saline or 10 HEPES, 70 mM NaCl, 0.5 mM EDTA, 5 M urea pH 7.4. The means of three experiments with standard deviation are plotted.

Most of the ^125^I-factor H and ^125^I-C1q remained in the fibrin clots after the urea wash. Thus, it is likely both proteins are covalently bound to fibrin. The covalent association of ^125^I-factor H and ^125^I-C1q still increased gradually with incubation beyond 180 min. When the fibrin clots were washed with urea, the binding of ^125^I-factor H and ^125^I-C1q was reduced. For example, the proportions of ^125^I-factor H and ^125^I-C1q binding were reduced from 22% to 13% and 13% to 11%, respectively at 30 min incubation. At 96 min incubation, 26% of factor H was shown to bind to fibrin clots after washing in non-denaturing buffer but only 18% remained bound after washing in urea buffer. Thus, non-covalently bound factor H or C1q were easily dissociated by urea. After 960 min incubation, approximately 32.5% of ^125^I-factor H and 17.6% of ^125^I-C1q remained bound to fibrin clots after washing in urea. Covalent binding between factor H and fibrin clots appeared to be complete at 960 min.

### Plasminogen interacts covalently with enhanced fibrin clots but HSA, Transferrin, IgG and α-2-Macroglobulin do not

Factor H and C1q are large asymmetric proteins which could possibly be trapped in the clots as fibrin crosslinking increases. To determine whether C1q and factor H interaction with fibrin clots is not simply by physical entanglement, binding assays was carried out using other plasma proteins with various molecular weights. These proteins were HSA (66 kDa), plasminogen (90 kDa), transferrin (81 kDa), IgG (150 kDa), α-2-Macroglobulin (720 kDa) as well as factor H (155 kDa) and C1q (410 kDa). These proteins were radioiodinated and added to human plasma which was supplemented with 2 mg/ml fibrinogen (final concentration). This formed enhanced fibrin clots by adding CaCl_2_ (final concentration of 20 mM) and incubation for 16 h at 37°C. Enhanced fibrin clots were washed three times in urea buffer and the amount of proteins bound to the fibrin clots was measured (Table 1). There was significant binding of C1q (28.5%), factor H (30%), and plasminogen (31%) to fibrin clots when 25,000 cpm was added to each reaction (0.5 nM factor H, 0.1 nM C1q, 1.4 nM plasminogen). Other plasma proteins (HSA, transferrin, IgG, α-2-Macroglobulin) showed low binding (< 5%) which could be the result of some traces of simple physical entrapment in the clots. Therefore, this suggests that factor H and C1q are retained in fibrin clots, not by physical entanglement but by specific binding followed by covalent crosslinking. Plasminogen was likely to be covalently bound to fibrin clots as the binding was stable after urea wash.

**Table 1:** Interaction of plasma proteins with fibrin clots. Enhanced fibrin clots were formed in the presence of various radiolabelled plasma proteins including ^125^I-factor H (5 ng), ^125^I-C1q (6 ng), ^125^I-human serum albumin (11 ng), ^125^I-plasminogen (12.5 ng), ^125^I-transferrin (20 ng), ^125^I-IgG (6.25 ng), and ^125^I-α-2-Macroglobulin (16 ng) (25,000 cpm/reaction used for each protein). The clotting reaction was allowed to proceed for 16 h at 37°C. After incubation, fibrin clots were washed three times in 10 mM HEPES, 70 mM NaCl, 0.5 mM EDTA, 5 M urea pH 7.4. The clot-bound proteins were measured as described above. The means of three experiments are presented with standard deviations.

| | Protein incorporated to fibrin clots (%±SD) |
| --- | --- |
| C1q | 30.2± 2.6 |
| Factor H | 28.7± 4.8 |
| Human Serum Albumin | 2.1 ± 1.0 |
| Plasminogen | 32.8 ± 5.6 |
| Transferrin | 1.7 ± 0.5 |
| IgG | 1.8 ± 0.4 |
| α-2-Macroglobulin | 2.3 ± 1.1 |

### Evidence for covalent binding of factor H and C1q to fibrin clots via FXIIIa (transglutaminase)

The reaction of ^125^I-factor H and ^125^I-C1q with fibrin clots was examined by SDS-PAGE. On a 4–12 % SDS-PAGE under reducing conditions, 5.2% of ^125^I-factor H (assessed by scanning the autoradiograph and analysis using Image Gauge software, Fuji FLA 3000 Imager) was observed as a large aggregate that did not migrate into the gel (Fig. 6A, *lane 5*). This suggests that the ^125^I-factor H was bound to fibrin clots covalently. There was also a small proportion of cross-linked products in the supernatant removed from clots (Fig. 6A, *lane 6*). When fibrin clots were formed with ^125^I-C1q and washed with urea, ^125^I-C1q was also found to be linked to the clots (Fig. 6B, *lane 5*). Therefore, ^125^I-C1q was also covalently bound to fibrin clots. It was of interest to see if this covalent binding involves crosslinking by FXIIIa. In order to investigate whether factor H and C1q are a target for FXIIIa, EACA or IAM were used as inhibitors. EACA, a competitive inhibitor for COOH and NH_2_ donor and acceptor groups, competes with fibrin for occupation of the enzyme active site. IAM inhibits the active site SH-group of FXIIIa and destroys its activity irreversibly. When the plasma was pretreated with EACA or IAM in the presence of ^125^I-factor H before clotting, no large aggregates were seen in both cases by SDS-PAGE and autoradiography (Fig. 6A, *lane3 and 4*). Similar results were obtained for C1q (Fig. 6B, *lane 3 and 4*). This suggests that factor H or C1q covalent binding diminished when FXIIIa was inhibited, either by EACA or IAM, confirming that the covalent interaction of ^125^I-factor H and ^125^I-C1q to fibrin clots occurs via FXIIIa.

**Figure 6.**
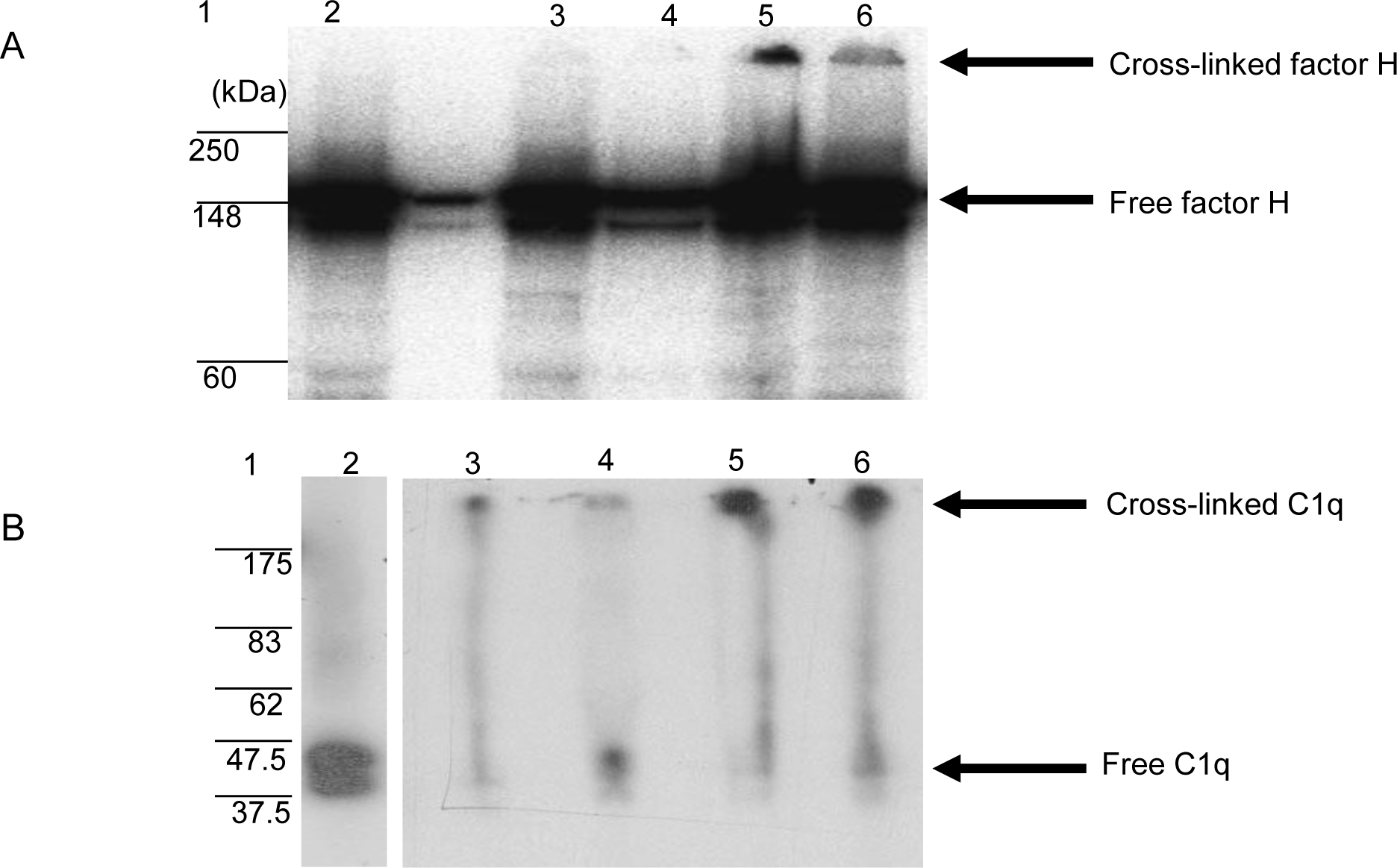
SDS-PAGE analysis of the ^125^I-factor H (A) and ^125^I-C1q (B) cross-linked to clots. Clots were formed in human plasma in the presence of ^125^I-factor H or ^125^I-C1q or ^125^I-plasminogen and the remaining bound materials were examined by SDS-PAGE (4-12% gradient gel) under reducing conditions. Various possible inhibitors of cross-linking, for example, epsilon amino caproic acid (EACA) or iodoacetamide (IAM), were incubated with plasma prior to addition of CaCl_2_. *Lane* 1, standard protein molecular mass marker; *lane* 2, standard ^125^I-factor H or ^125^I-C1q; *lane 3*, incubation with EACA before Ca^2+^; *lane 4*, addition of IAM before Ca^2+^; *lane 5*, ^125^I-factor H or ^125^I-C1q bound to clots without inhibitors; lane 6, ^125^I-factor H or ^125^I-C1q from the supernatant of the clotting mixture. It is likely that ^125^I-factor H and ^125^I-C1q bound to clots shows up as a band corresponding to the cross-linked products which remained in the well (*lane 5*). No cross-linked products were observed with inhibitor controls (*lanes* 3 and 4).

### Complement classical pathway is activated by fibrin clot formation

Since factor H and C1q both bound non-covalently and covalently to clots formed in plasma, it was of interest to examine whether clots activated the complement system as would be expected from the binding of C1q, and also, whether factor H regulated complement activation induced by fibrin clots. To examine the effects of clot formation on complement activation, a haemolytic assay for C4 was carried out with sensitized SRBC and either normal or factor H-depleted plasmas (in the presence or absence of clotting). This assay, using C4 deficient guinea pig serum, measures the relative amount of classical pathway activation as determined by C4 consumption. The C4 assay was used because this is specific for classical and (lectin) pathway and to avoid involvement of C3, as factor H-depleted plasma becomes secondarily depleted of C3. When serially diluted normal and factor H-depleted plasmas were incubated with sensitized EA and C4 deficient guinea pig serum, comparison between normal and factor H-depleted plasmas showed that there was a small loss of C4 activity in the factor H-depleted plasma through factor H-depletion procedures (Fig. 7). In order to assess the complement activation, possibly induced by clot formation, fibrin clots were initially formed in the presence or absence of factor H using normal or factor H-depleted plasmas. After clotting, supernatants were assayed for C4 activity. The results demonstrated that in normal plasma after clotting, there was a decrease in C4 activity by 42.5% compared to normal unclotted plasma. In factor H-depleted plasma there was a 65.3% decrease in C4 activity after clotting compared to factor H-depleted unclotted plasma (Fig. 7). Thus, clots activate the classical pathway of complement in normal and factor H-depleted plasma. Moreover, C4 depletion induced by clotting of the factor H-depleted plasma was approximately 50% greater than that in normal plasma.

**Figure 7:**
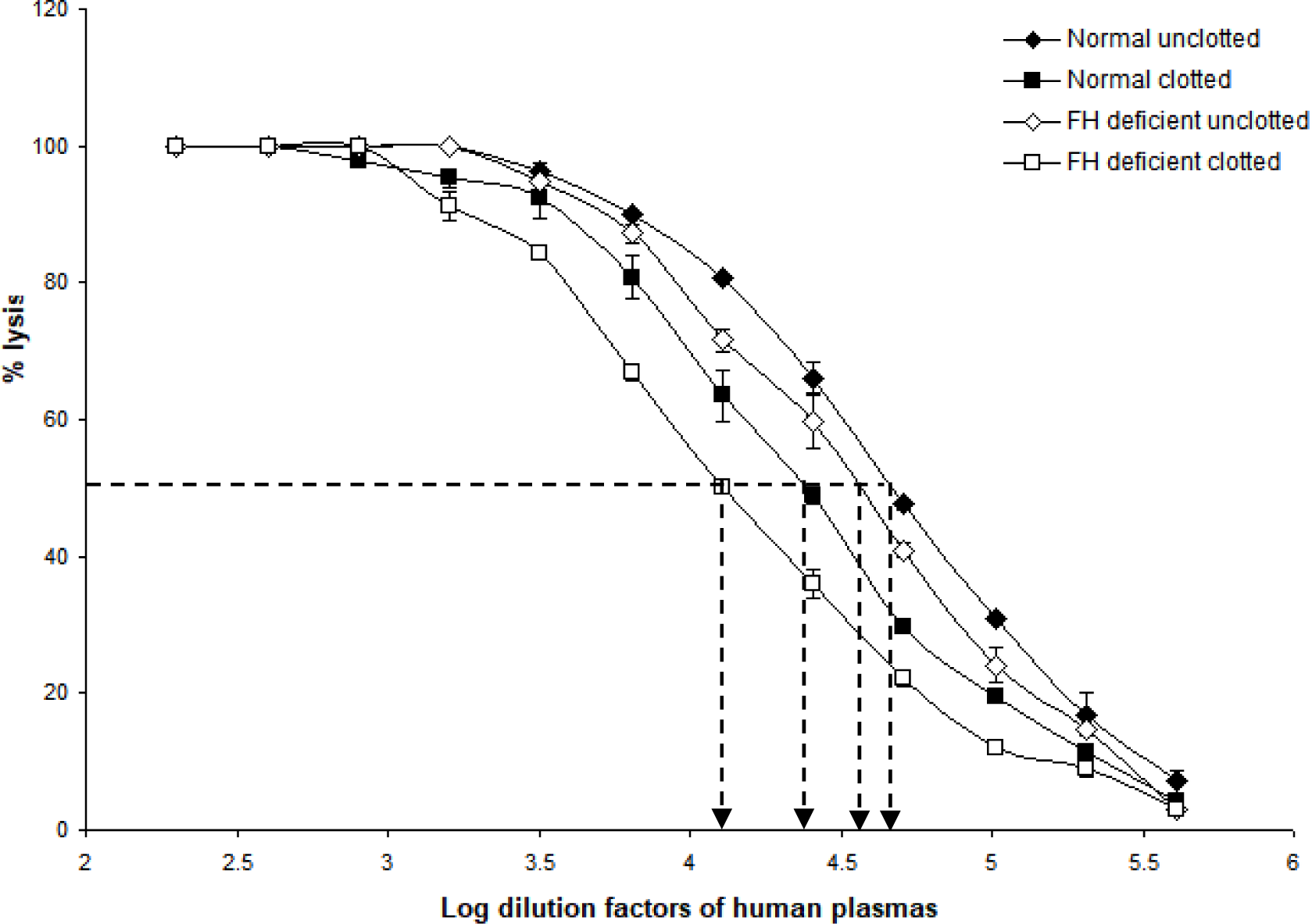
Activation of complement by clot formation in the absence of factor H. Clots were formed in normal and factor H-depleted human plasmas and supernatants of each sample were tested to assess the remaining complement C4 activity. Supernatants removed from normal and factor H-depleted human plasmas (1/200 up to 1/409,600) were serially diluted in DGVB^2+^ and each dilution was incubated with 1/32 dilution of C4-deficient guinea pig serum and sensitized sheep red blood cells (EA) for 1 h at 37°C. Normal and factor H-depleted plasmas (dilution of 1/200 to 1/409,600) without clotting were also tested for C4 activity. Experiments were performed in duplicate and the average is shown. C4 consumption by clot-induced complement activation was calculated as follow. The dilutions of plasmas required to provide 50 % lysis were read off from the graph. C4 activity is expressed as the reciprocal dilution of serum/plasma required to give 50% lysis. It expressed as units of C4 activity per unit volume so for example, normal human plasma gives 50% lysis at 1:41687 dilution, and can be said to have 41687 activity units of C4 per unit volume. The amount of C4 consumption by clots were calculated by comparing with unclotted plasma (normal human plasma vs normal human plasma after clotting; factor H-depleted plasma versus factor H-depleted plasma after clotting). The results demonstrated that in normal plasma after clotting, there was a decrease in C4 activity by 42.5% compared to normal unclotted plasma. In factor H-depleted plasma, there was a 65.3% decrease in C4 activity after clotting compared to factor H-depleted unclotted plasma.

Complement activation by fibrin clot formation was also measured by MAC formation as this determines whether the classical pathway of complement is fully activated by clots i.e. up to the C9 stage. This was performed by incubating fibrin-coated wells with serially diluted fresh human serum. MAC deposited on the fibrin was then measured directly from the fibrin-coated well after removing the human serum. Incubation of the same dilutions of human serum with uncoated wells (blocked with PBS-0.5 mM, 0.1% Tween 20) was used to measure non-specific formation of MAC in the well. This was then subtracted from each sample value. As shown in Fig. 8A, there was detectable MAC on the fibrin-coated wells when the wells were incubated with 1/10, 1/50, 1/250 diluted human serum. Fibrinogen-coated wells showed lower MAC deposition with 1/10, and 1/50 dilutions of human serum and ovalbumin-coated wells also showed a low level of deposition. Ovalbumin antigen-antibody complex-coated (OVA ag-ab) wells were used as a positive control and showed a high level of MAC formation compared to fibrin- or fibrinogen-coated wells. This suggests that fibrin activated complement which led to formation of MAC. Interestingly, MAC was detected on the fibrin-coated wells, showing that MAC could also be bound on the surfaces of fibrin clots. MAC was also directly detected on OVA ag-ab complex-coated wells. When MAC is formed, it is inserted in the lipid bilayer of the plasma membrane of cellular complement activators. Otherwise, it can react with any of several plasma proteins, for example clusterin or S-protein (vitronectin), which prevent insertion into lipid bilayers. Since the wells used in this system have no lipid bilayer, the MAC must be binding to some structure on the fibrin or OVA ag-ab complexes. The MAC-plasma protein complexes (for example, SC5b-9, the complex with S-protein) might also bind to these surfaces. The assays used do not distinguish between MAC (C5b-9) and the other forms such as SC5b-9. In a capture ELISA system (Fig. 8B), MAC was measured from each supernatant of serially diluted human serum that was incubated with fibrin-coated wells. There was a high level of MAC in the supernatants from fibrin-coated wells, a lower level from fibrinogen-coated wells, and a low level from OVA ag-ab complex-coated wells. This suggests that both fibrin and immune complexes activate complement to produce MAC. However, for the immune complexes, most of the MAC binds to the immune complexes, while for fibrin, most of the MAC remains in solution. This confirms that loss of C4 activity on clotting is due to complement activation, and not due to sequestration of unactivated C4 by the clots.

**Figure 8:**
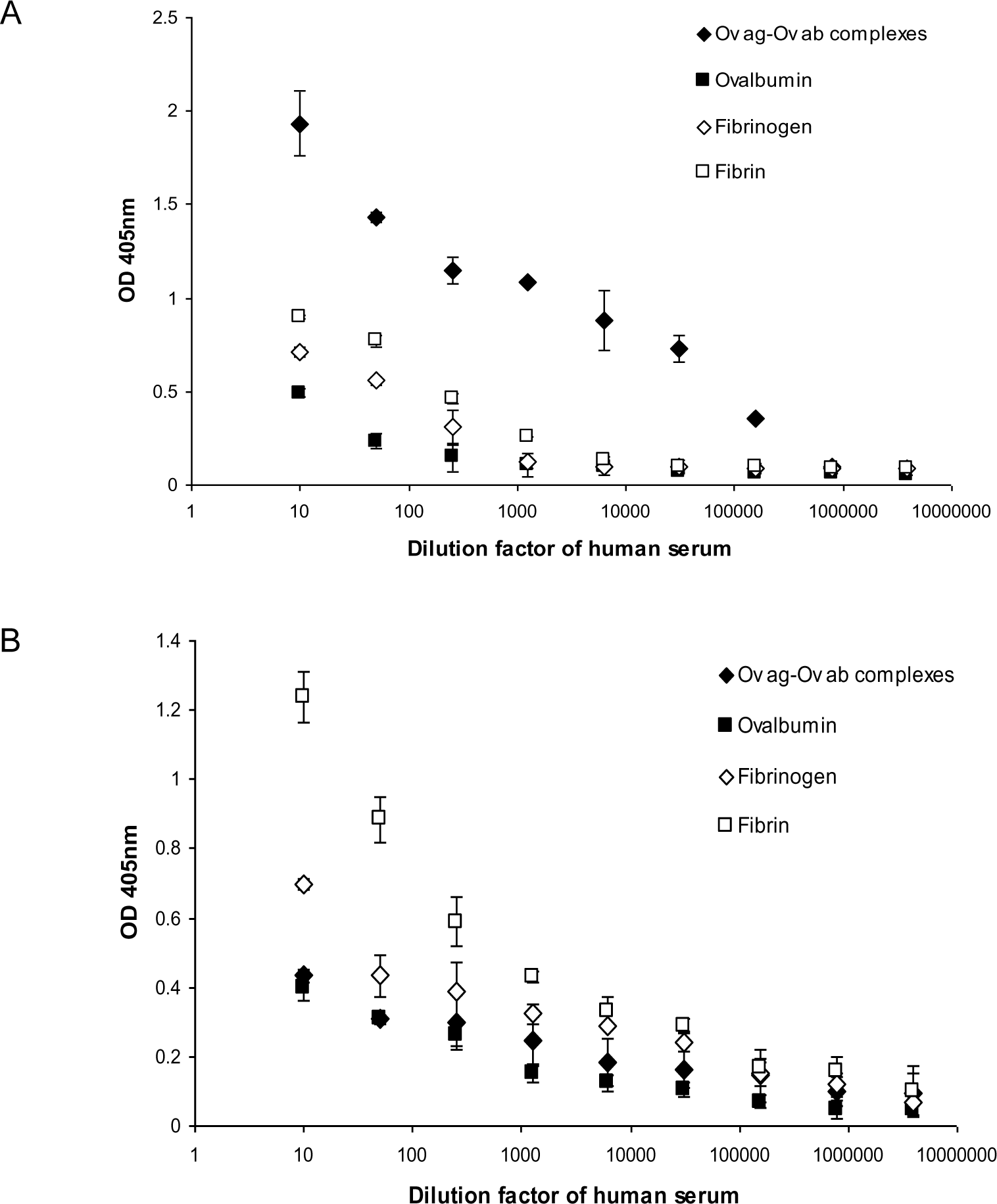
Detection of membrane attack complex (MAC) on fibrin-coated wells (A) and in the supernatants of human serum incubated with the fibrin-coated wells (B). (A) Serial dilutions of fresh human serum (1/10, 1/50, 1/250, 1/1250, 1/6250, 1/31250, 1/156250, 1/781250, 1/3906250) were incubated with ovalbumin antigen-antibody complexes-, ovalbumin-, fibrinogen- and fibrin-coated wells. After incubation, the supernatant from each well was removed and the well was washed three times in PBS-0.1% Tween20. MAC was detected in ovalbumin antigen-antibody complexes-, ovalbumin-, fibrinogen- and fibrin-coated wells with mouse anti-neo C9 antibody using an ELISA system. (B) MAC was also detected in the supernatants in a Capture ELISA system with rabbit anti-C9 monoclonal antibody. To measure the non-specific formation of MAC, serial dilutions of fresh human serum were incubated with blocked wells alone, and MAC from each well and the supernatant were assayed in the same ELISA and Capture ELISA systems as above. These values were subtracted from each sample as a background. The means of three experiments are presented with standard deviations.

### C1q and factor H bind to enhanced clots formed in the presence and absence of plasma

It was further investigated whether the binding of factor H and C1q to fibrin clots is direct or whether it could be mediated via other plasma proteins. The assays used here were fibrin immobilised on microtitre plates and fibrin clots formed in human plasma. However, although we were able to show that factor H binds to fibrin-coated wells, binding of C1q to fibrin-coated wells was uncertain as it was inhibited by BSA. Moreover, binding of factor H and C1q to clots formed in plasma was high but it was not certain whether factor H and C1q bind directly to fibrin or to other proteins present in the plasma clots. Therefore, it was necessary to examine the binding to fibrin-only-clots. Thus, to consolidate this work, fibrin-only-clots and as a control, enhanced plasma clots were used. The results show that binding of ^125^I-factor H to enhanced plasma clots and fibrin-only-clots was 21.5% and 19%, respectively, when the clots were washed in HEPES-½-saline (Fig. 9). Binding of ^125^I-C1q to enhanced plasma clots and fibrin-only-clots was also seen and the amount of ^125^I-C1q bound to both clots was very similar. When the clots were washed in urea buffer, the binding of both ^125^I-factor H and ^125^I-C1q to enhanced plasma clots and fibrin-only-clots was reduced compared to the clots washed in HEPES-½-saline, but for both types of clots, there was still substantial binding of both factor H and C1q. The binding of factor H and C1q to the fibrin in the clots is therefore mainly direct, and does not require the presence of other plasma proteins.

**Figure 9:**
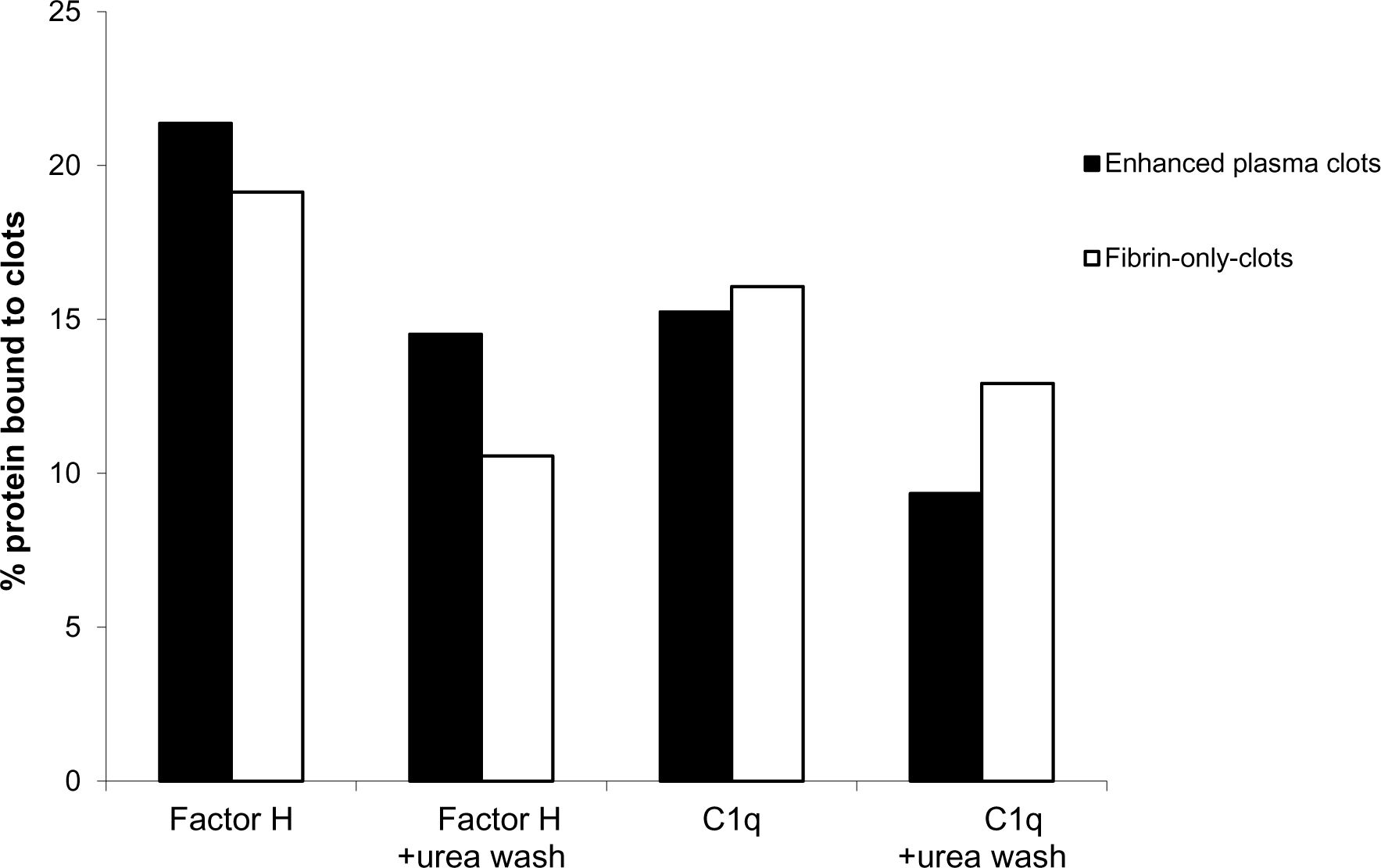
Binding of ^125^I-factor H or ^125^I-C1q to “enhanced clots” formed in the presence or absence of plasma. Enhanced plasma clots were formed in the presence of ^125^I-factor H or ^125^I-C1q (25,000 cpm/reaction). Clots were washed three times with 500 μl of HEPES-½-saline buffer or 10 mM HEPES, 70 mM NaCl, 0.5 mM EDTA, 5 M urea pH 7.4 The concentration of fibrinogen in the “enhanced clot” mixture is 3.5 mg/ml. To make clots in the absence of plasma, 3.5 mg/ml purified fibrinogen in HEPES-½-saline buffer was used. The fibrinogen solution (10 μl of 3.5 mg/ml) was premixed with ^125^I-factor H or ^125^I-C1q (25,000 cpm/reaction) in a total volume of 96.5 μl in 10 mM HEPES, 70 mM NaCl, 5 mM CaCl_2_ pH 7.4. Clotting was initiated by adding 3.5 μl of thrombin (8.75 μg/ml, final concentration). This quantity of thrombin was selected as incubation of fibrinogen with thrombin at the same weight ratio resulted in complete cleavage of the α- and β-chains of fibrinogen at 40 min incubation at 37°C, as judged by SDS-PAGE analysis in Fig. 6. As for the enhanced clots, the mixture was incubated for 40 min at 37°C and the resulting clots processed and washed as for the “enhanced clots”. ^125^I-factor H or ^125^I-C1q which remained associated with clots was measured.

### Reduction of factor H and C1q levels in human serum compared to those in plasma

In order to examine whether factor H or C1q is bound to clots, factor H or C1q level in plasma and the serum from the same plasma, was assayed in an ELISA system. Five samples of citrated plasmas with different ages in storage and conditions were tested (Fig. 10). The results showed that in all five samples, both factor H and C1q levels were reduced in sera compared to the plasmas. In a healthy volunteer sample, approximately 38% of factor H level was reduced relative to the plasma and 15 % of C1q level was lost in serum compared to the plasma. Thus, our results support previous finding that factor H and C1q bind to fibrin clots.

**Figure 10:**
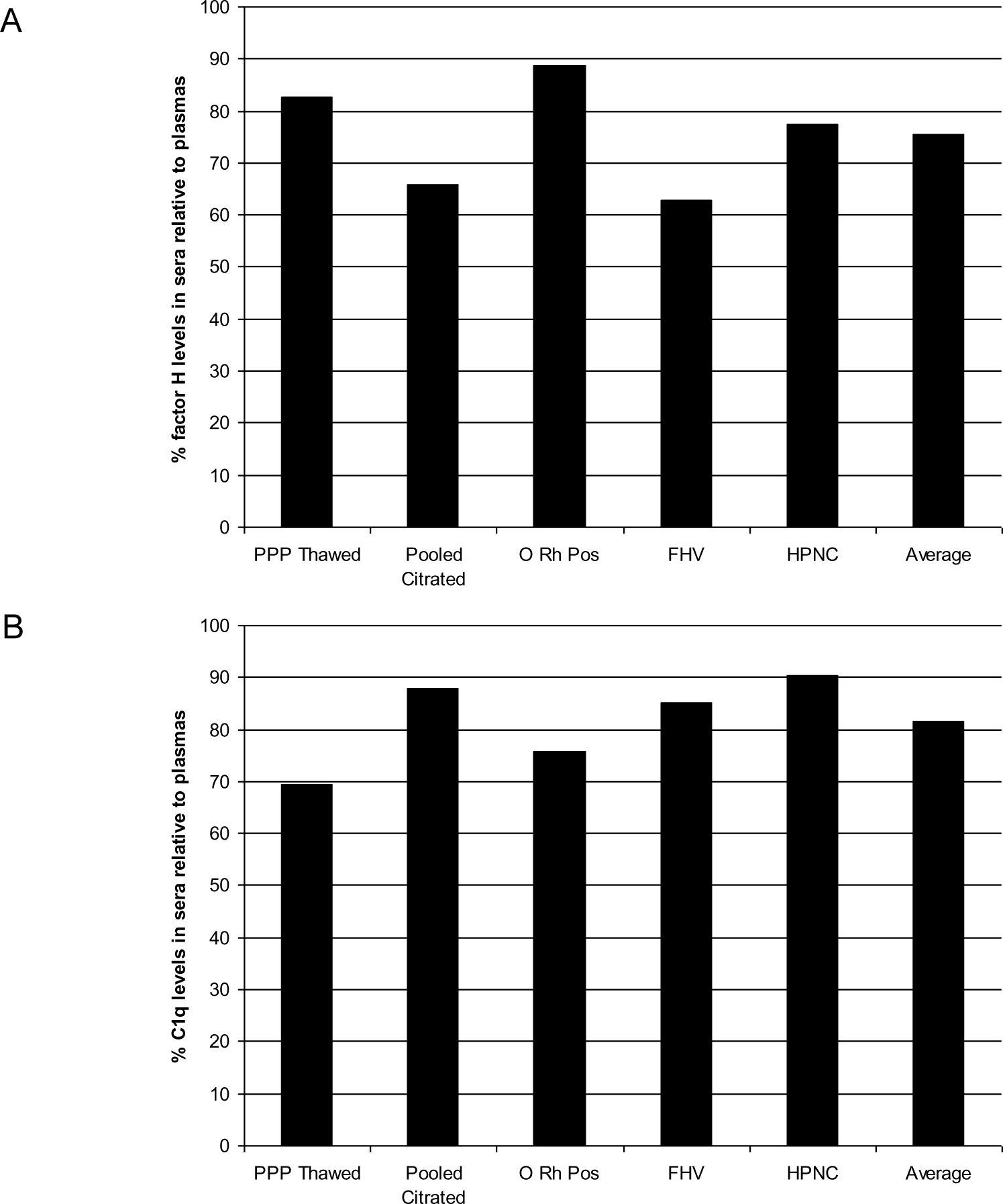
Measurement of relative factor H or C1q levels in human sera compared to those in plasmas. In order to investigate whether factor H or C1q is bound to clots, factor H or C1q levels in plasma and the serum from the same plasma, were assayed. Five samples of citrated plasmas with different ages in storage and conditions were tested. Serial two-fold dilutions of each plasma and the serum (1/20 to 1/10240) were incubated with monoclonal anti-human factor H antibody-coated wells for 1 h at 20°C. After incubation, the plates were washed in PBS-0.1% Tween20 and factor H was detected with rabbit-human factor H using an ELISA system. C1q was also measured from the same dilutions of the plasma and the serum used in the factor H ELISA. Each dilution was incubated with ovalbumin antigen-antibody complexes-coated wells for 1 h at 20°C. C1q was detected with biotinylated rabbit anti-human C1q antibody. For each sample, a graph of dilution factors of the plasma and the serum against OD_405nm_ were drawn. The dilution factors of the plasma and the serum at OD_405nm_ of 2.5 were read off from the graph. The percentage of the factor H level in the serum was calculated by comparing with that in the plasma (assuming that the factor H level in the plasma was 100%). The percentage of the C1q level in each serum sample was also calculated as same as above. Average represents the mean of relative factor H or C1q levels of five sera samples. PPP Thawed, O Rh Pos, FHV and HPNC mean platelet-poor plasma, O Rhesus positive plasma, fresh plasma from a healthy volunteer and human normal plasma citrated, respectively.

## DISCUSSION

### Factor H and C1q interact with fibrin clots covalently via FXIIIa

It is generally believed that coagulation activates complement in mouse blood. Because it is very difficult to conserve complement activity in mouse serum, anticoagulated plasma is nearly always used for measurement of mouse complement activity. Here, we asked whether clotting affects complement activity in humans. Both factor H and C1q are reported to recognise “foreign” and altered-self materials. It was therefore of interest to ascertain whether factor H has further associations with clots since clots are an example of “altered-self”. As a further connection between complement and coagulation, factor H is present in platelets. It might therefore be possible that when clotting or clots made of fibrin or fibrin and platelets, stimulate complement activation, factor H is present to downregulate this process.

We investigated whether fibrin, which is the main protein constituent of blood clots, activates complement. Therefore, we examined whether factor H interacts with fibrin clots. The experiments were performed in two stages, using first a microtitre plate assay and later in a more physiological clotting assay. The results obtained in the plate assays demonstrate that factor H and C1q interact with fibrin immobilised on plates, and for both, binding to fibrin is greater than to fibrinogen (Fig. 1). The specificity of C1q binding was uncertain as BSA inhibited C1q binding. Factor H, however, binds to fibrin clots with minimal inhibition by BSA. Factor H binding was shown to be dose-dependent and saturable (Fig. 2). Scatchard plots further show that the affinity of factor H binding for fibrin is exceptionally high (Kd 55 pM), and there is one major class of binding sites on fibrin-coated wells. Ionic interactions are important for factor H-fibrin binding, as there was a decrease in interaction with an increase in salt strength. The quantity of factor H and C1q bound to fibrin plates was, however, very small. This could be due to limited number of binding sites present in fibrin-coated wells. It was difficult to envisage how fibrin could be made to bind to plates in a correct physiological configuration (i.e. polymerised), so the presentation of fibrin on the plates used was likely to be very artificial. However, the apparent specificity of factor H binding and preference for fibrin over fibrinogen justified continuing the investigation using a system closer to physiological conditions. Clotting of whole plasma or of plasma supplemented with extra fibrinogen was used. The results indicated that factor H and C1q do bind to clots, and that a large proportion of the bound factor H and C1q may be covalently bound (Fig. 4 and 5). The maximal binding of factor H and C1q to fibrin clots was very high (i.e. a high percentage of the total factor H and C1q present in plasma), in contrast to the results with the plates. There was evidence of cross-linking taking place between fibrin monomers for a period of time as clot sizes decreased after 96 min incubation and further reduction in size (approximately 10-fold) was observed after 16 h incubation. FXIIIa is a critical component in the coagulation pathway which cross-links adjacent COOH and NH_2_ groups in neighbouring fibrin molecules (62, 63). It was tested whether factor H becomes covalently associated with fibrin clots using a urea wash assay of the clots and SDS-PAGE analysis. In the clot–urea wash assays, clots were washed firstly with non-denaturing buffer, and secondly, with denaturant containing urea. At 96 min incubation, approximately 18% factor H remained bound to clots after washing in 5 M urea compared with 25% after washing in non-denaturing buffer (Fig. 5). Binding of factor H to clots increased up to 16 h incubation at which time most bound factor H molecules were no longer dissociated by urea i.e. were covalently linked to clots. The binding of C1q was approximately 24% (after 16 h incubation) when the clots were washed with non-denaturing buffer, but only 18% of bound C1q remained associated with clots after washing in denaturant for the same incubation time. Although C1q binding was partially dissociable with denaturant at 16 h incubation, the result indicates that both factor H and C1q binding is covalent. SDS-PAGE analysis (Fig. 6) showed that ^125^I-factor H and ^125^I-C1q were incorporated into a large aggregate, which is likely to be factor H and C1q cross-linked to clots. This evidence was supported by control experiments where no high molecular weight aggregates were seen when FXIIIa was inhibited by either EACA or IAM. Therefore, we can conclude that FXIIIa crosslinks factor H and C1q into clots.

FXIII consists of two catalytic A subunits (FXIIIA) and two noncatalytic B subunits (FXIIIB) held together by noncovalent bonds. Human FXIIIB is composed of 10 CCP domains which are the characteristic domains of the regulatory proteins of complement activation system including factor H. Moreover, a significant structural similarity between FXIIIB and factor H was shown with a high degree of amino acid identity between the FXIIIB CCP5 and factor H CCP16 and 18 (64). Thus, it is likely that factor H present in the platelets interacts with FXIIIa which then crosslinks into the clots. The proportion of factor H which is covalently bound, is very high (approximately 32%) in the clot-urea wash assay (Fig. 5). In the SDS-PAGE analysis, however, only 5% of the radioactivity is in the large aggregate (Fig. 6). There are several possibilities for the quantitative differences in cross-linked factor H analysed in the clot-urea wash assay and the SDS-PAGE. Firstly, in the clot-urea wash assay, the amount and concentration of plasma used were larger than that in the SDS-PAGE analysis (i.e. 45% plasma, final concentration in the final volume of 100 μl) was used in the clot-urea wash assay whereas only 0.9% plasma (final concentration in the final volume of 50 μl) was used in the SDS-PAGE). Secondly, the amount of extra fibrinogen used in the clot-urea wash assay was higher (200 μg in 100 μl) than that in SDS-PAGE (20 μg in 50 μl). Thirdly, SDS-PAGE analysis may underestimate the amount of factor H covalently bound. Because the factor H cross-linked to clots remained in the loading well, there may have been considerable loss of crosslinked factor H while processing the gel after electrophoresis. However, the results obtained from both assays suggest that factor H binds to clots covalently and the percentage of factor H covalently linked to clots is high: 5% (obtained from SDS-PAGE) - 32% (obtained from the clot-urea wash assay). The phenomenon of covalent binding may account for the unusually high affinity of factor H for fibrin on the coated plates (Fig. 5). The very low Kd could be an artefact generated by some proportional of factor H becoming covalently bound, since FXIIIa will also be on the plates just as it is on the clots.

In order to establish that factor H and C1q binding is not a simple entanglement, several plasma proteins were assessed for binding to fibrin clots (Table 1). Only ^125^I-plasminogen was shown to be bound to clots covalently. This was expected because fibrin is the major plasmin substrate. Plasminogen can bind to fibrin though lysine residues and it can be converted to plasmin by tissue plasminogen activator as well as by urokinase (21). However, other plasma proteins (HSA, transferrin, α2Macroglobulin, IgG) did not interact with clots. These proteins span the size range of C1q and factor H. This suggests that binding of factor H, C1q and plasminogen is specific and it is not a simple physical entanglement. SDS-PAGE analysis (Fig. 6) further showed that FXIIIa is essential for covalent binding of factor H and C1q to clots.

The binding of both factor H and C1q is low to fibrin-coated plates but very high to clots. Explanations for the differences are likely to be: (i) fibrin clots made on plates are “artificial” and randomly oriented. It was difficult to coat the plates with fibrin directly so that the fibrin clots were made on the plates by adding fibrinogen and thrombin. Addition of fibrinogen onto the plates prior to thrombin, can limit the mobility of fibrinogen necessary for its cleavage by thrombin. This can produce an unknown extent of conversion of fibrinogen to fibrin. (ii) In clots, other proteins may be present which mediate binding and thus it is possible that binding may not be to fibrin. As shown in Fig. 9, however, binding of factor H and C1q to fibrin in clots is mainly direct. (iii) The surface area of the clots is likely to be very large compared with the plate but this is difficult to measure. The quantity of fibrin in clots was much greater than on plates. (iv) In order to optimise clot formation and potential binding to fibrin clots by ^125^I-factor H or ^125^I-C1q, fibrinogen (2 mg/ml) was added to a reaction mixture prior to clot formation (referred to “enhanced clots”). Thus, this increases the surface area of clots for factor H or C1q binding which in turn, increases the concentration of “epitopes” for factor H binding. (v) Covalent-linking of factor H and C1q in clots by FXIIIa (the concentration of FXIII in plasma is 10 μg/ml) makes binding irreversible, so that no dissociation can occur, therefore, apparent avidity of factor H or C1q is increased.

### Factor H regulates classical complement pathway activation at the site of clotting?

Previous studies have established that factor H can regulate the classical pathway activation triggered by certain targets (11–13). Notably, factor H has been found to compete with C1q for binding to cardiolipin, a known activator of the classical pathway, thereby preventing its activation (13). Thus, Factor H can act as a control switch for C1q-mediated complement activation, thereby preserving immune homeostasis. Similarly, C1q binding to lipid A, a component of the outer membrane of Gram-negative bacteria, typically initiates the classical pathway activation (12). However, factor H can outcompete C1q in this context too, effectively regulating the immune response towards non-self-entities like bacteria. Moreover, factor H also has implications in the clearance of apoptotic cells, or dying cells, by phagocytes, a process that is typically enhanced by C1q (11, 65). It appears that factor H can modulate the C1q-enhanced uptake of apoptotic cells, possibly by acting as a bridge between apoptotic cells and phagocytes. This role of factor H could potentially modify signalling pathways to reduce the pro-inflammatory effects often associated with the process, adding yet another layer of immune regulation. By outcompeting C1q for binding to both self and non-self-ligands and controlling phagocytosis of apoptotic cells, factor H helps maintain a balanced immune response and protects against potential self-damage.

Here, we wished to investigate complement classical pathway activation by clots, and a possible role for factor H in downregulating this. In order to measure the classical pathway activation, a C4 consumption assay was performed using normal human plasma or factor H-depleted plasma. The results indicate that clots activate complement both in normal and factor H-depleted plasma (Fig. 7). In addition, assays for MAC deposition also demonstrated that complement was activated by clots fully up to C9 stage and the reaction was completed by the formation of MAC (Fig. 8). Thus, C4 consumption by clots was due to C4 activation, not just C4 binding to the clots. Moreover, there is an increase in C4 consumption by clots in the factor H-depleted plasma compared to the normal plasma. This suggests that in the absence of factor H, C4 consumption by the classical pathway, triggered by clots is higher by 50 % than in normal plasma. These results support the hypothesis that factor H downregulates the classical pathway activation, possibly by competing with C1q for binding to fibrin clots. However, we were unable to demonstrate direct competition between C1q and factor H in this clotting assay because the concentration of factor H present in the plasma is high (between 150-750 μg/ml) so that it was not practical to compete out factor H binding to fibrin clots by C1q. Moreover, we could not perform an assay to measure direct competition between C1q and factor H to fibrin using plate binding assay. This was because specificity of C1q binding to fibrin-coated wells was not certain as the binding was inhibited by BSA. Further, factor H binding to fibrin coated on microtitre plates was only detectable at low salt strengths (20 mM NaCl), and at this salt strength, we found that factor H and C1q interact with each other (data not shown).

These results may eventually further our understanding of diseases, where the complement and the coagulation systems play a crucial role in the pathology such as CoVID-19. The severe acute respiratory syndrome coronavirus 2 (SARS-CoV-2) infection, responsible for the CoVID-19 pandemic, has been associated with a multisystem and multiorgan inflammatory response that contributes to high mortality rates in affected individuals (66). Central to this response is the hyperactivation of the complement and coagulation systems (67–69). Serum samples from CoVID-19 patients have revealed a consistent and sustained activation of complement system. It has been found to play a significant role in the hyper-inflammation and thrombotic microangiopathy observed in severe CoVID-19 cases (66). SARS-CoV-2 has the ability to trigger the activation of all three pathways of the complement system: the classical, alternative, and lectin pathways (70–72). Furthermore, SARS-CoV-2 was also found to promote the synthesis and release of complement factors from infected respiratory epithelial and endothelial cells through Janus kinase 1 (JAK1)-dependent and/or JAK2-dependent pathways, interestingly parallel to the release of procoagulant (clot-promoting) factors (73, 74). The activation of the complement system, specifically the cleavage of C5, can result in the release of prothrombotic factors (74). Additionally, C5 cleavage can increase the expression of P-selectin on endothelial cells and trigger tissue factor (TF) activation, both of which facilitate the recruitment of inflammatory leukocytes (74). TF can be expressed on Neutrophil Extracellular Traps (NETs) which are released by neutrophils during an immune response (67). This expression of TF can contribute to clotting, a condition that is often associated with severe cases of CoVID-19 (75, 76). This is one of the mechanisms by which a hyperactivated complement system could initiate coagulation or blood clotting. Furthermore, TF activation has been noted in the presence of SARS-CoV-2, and there is evidence that TF expression can be disrupted by inhibiting the complement component C3, suggesting that the complement axis plays a critical role in CoVID-19 induced inflammation (77). The coagulation system also plays a pivotal role in the pathology of COVID-19. The interaction of the spike glycoproteins of SARS-CoV-2 with ACE2 receptors on the host cell surface leads to injury in the vascular wall of blood vessels, leading to coagulation and clotting cascades activation and subsequent formation of internal blood clots (67). Furthermore, severe CoVID-19 is associated with coagulopathies exacerbated by the formation of NETs, which can block pulmonary lymphatic vessels (78). These clots are often associated with systemic inflammation, disease severity, comorbidities, and increased mortality risk (79). Thus, the activation of the complement system by SARS-CoV-2, along with the formation of clots, presents a significant risk factor in severe CoVID-19 cases. Hence the reported interactions between the complement and coagulation systems, and its dysfunction could play a crucial role in the hyperinflammation and thrombotic complications observed in severe CoVID-19 cases. However, further research is needed to fully elucidate these mechanisms and their implications for CoVID-19 pathology and treatment.

## CONCLUSION

This study provides significant insights into the complex interplay between the coagulation and complement pathways, specifically the interaction between factor H, C1q, and fibrin clots (Fig. 11). Our results indicate that factor H and C1q specifically bind to fibrin clots, with a considerable proportion being covalently bound. Moreover, it was found that clots activated the classical pathway, and factor H can restrict this activation, supporting the idea of factor H having a regulatory role in the classical pathway. The work also suggests that FXIIIa, a critical component in the coagulation pathway, crosslinks factor H and C1q into clots. These findings offer a new understanding of how these molecules interact during coagulation and complement activation, which could be pivotal in pathophysiological conditions where these processes are dysregulated, such as CoVID-19, and hence, raises the possibility of therapeutic interventions targeting these molecular relationships.

**Figure 11:**
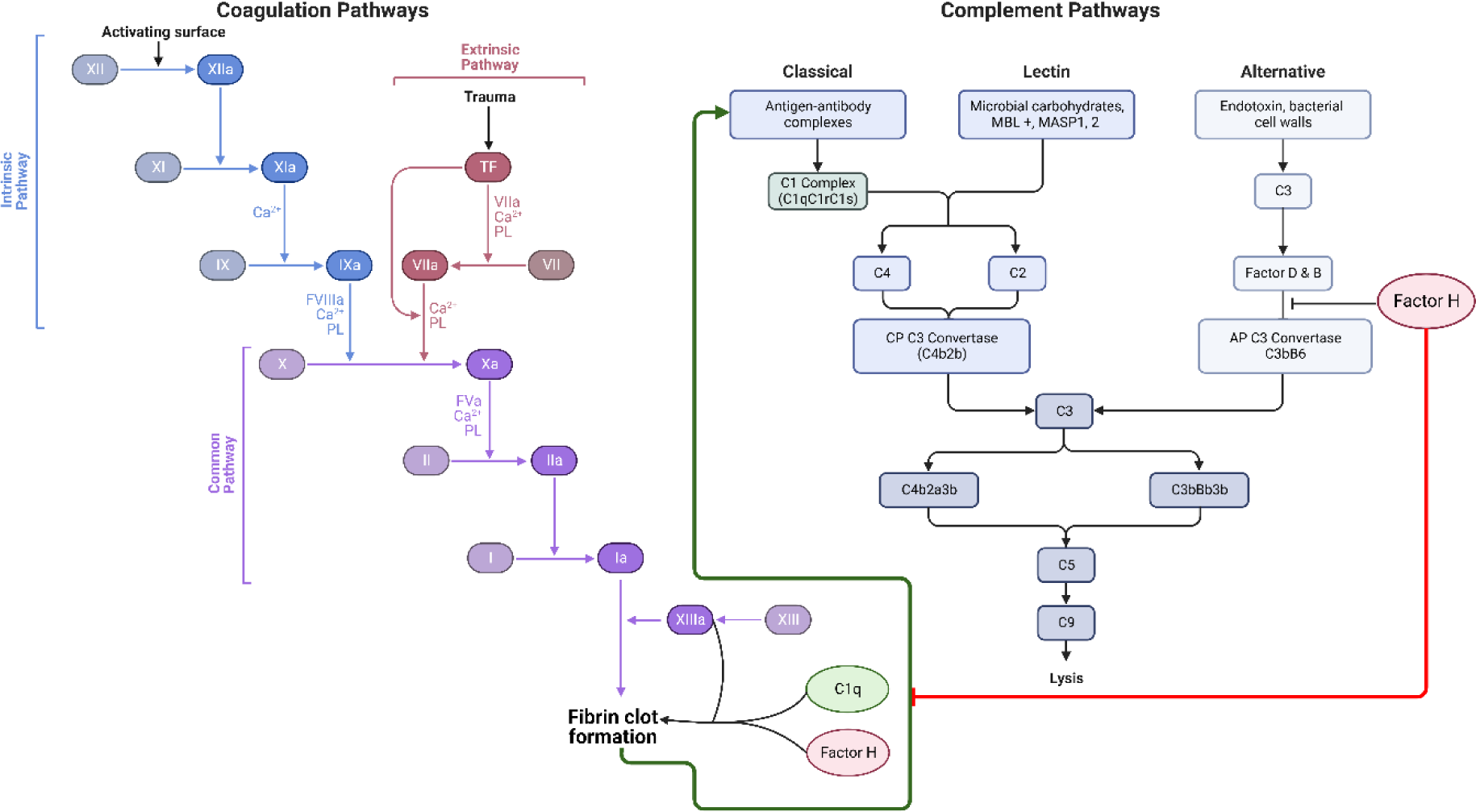
Novel cross talk between the coagulation and complement system. This graphic abstract summarises the distinct yet interconnected mechanisms of the complement and coagulation pathways. The complement classical pathway is initiated by the binding of C1q to IgG or IgM-containing immune complexes or other non-immunoglobulin targets, while the lectin pathway is activated when mannan-binding lectin (MBL) encounters conserved pathogenic carbohydrate motifs, leading to the activation of the MBL-associated serine proteases (MASPs). Activation of the classical or lectin pathways results in the cleavage of C4 and C2, producing the C3 convertase (C4b2a), which cleaves C3 to form C3b. In the alternative pathway, spontaneous hydrolysis of C3 yields a C3b-like molecule [C3(H_2_O)], which binds to Factor B, facilitating its cleavage by Factor D to produce Bb. The resultant C3(H_2_O)Bb, analogous to C3 convertase C4b2a, cleaves C3 to generate C3b. Covalently bound C3b on target surfaces forms more convertase, C3bBb, and the association of C3b with either C4b2a or C3bBb transforms them into classical or alternative pathway C5 convertases, respectively. The cleavage of C5 by these convertases initiates the assembly of the Membrane Attack Complex (MAC; C5b-C9), which binds to microbial surfaces, potentially leading to lipid bilayer membrane lysis. The coagulation pathway is initiated by vascular wall damage and is subdivided into intrinsic, extrinsic, and common pathways. The extrinsic pathway is triggered by tissue factor exposure, activating factor VIIa and calcium. The intrinsic pathway starts with factor XII exposure to subendothelial collagen, setting off an activation cascade involving factors XI, IX, and VIII. Both pathways culminate in factor X activation, marking the onset of the common pathway. Factor X, alongside factor V, forms the prothrombinase complex, catalyzing the conversion of prothrombin into thrombin, which then mediates the polymerization of fibrinogen into fibrin monomers, cross-linked by factor XIII to form a stable fibrin clot, which is able to capture platelets and red blood cells, effectively sealing the wound and stemming plasma loss. The coagulation cascade and the complement system communicate through many direct and bidirectional interactions. A novel interaction described in this study is between factor H, C1q, and fibrin clots. Factor H and C1q have been shown to covalently bind to fibrin clots with high specificity, and FXIIIa aids in cross-linking factor H and C1q into the clots. Moreover, clots activate the complement classical pathway, with factor H dampening this activation, suggesting a regulatory role for factor H in the clot-induced activation of the classical pathway. The figure was created using templates provided at Biorender.com.

